# Targeting epilepsy with photopharmacology in human brain tissue

**DOI:** 10.64898/2026.02.13.705713

**Authors:** Mayan Baues, Ahmed Elgokha, Mai My Hong Nguyen, Merlin Schwering-Sohnrey, Tongil Ko, Florian Schwermer, Filomena Perri, Thoralf Opitz, Tony Kelly, Gilles Huberfeld, Valeri Borger, Matthias Schneider, Rainer Surges, Heinz Beck, Dirk Trauner, Christa E. Müller, Michael Wenzel

**Affiliations:** Department of Epileptology, University Hospital Bonn, Bonn, Germany; Institute of Experimental Epileptology and Cognition Research, Medical Faculty, University of Bonn, Bonn, Germany; Department of Pharmaceutical & Medicinal Chemistry, Pharmaceutical Institute, University of Bonn, Bonn, Germany; Department of Chemistry and Department of Systems Pharmacology and Translational Therapeutics, University of Pennsylvania, Philadelphia, USA; Université Paris Cité, Institute of Psychiatry and Neuroscience of Paris (IPNP), INSERM U1266, Neuronal and Astroglial Signaling in Epilepsy and Glioma, 75014 Paris, France; Service de Neurologie, Hôpital Fondation Adolphe de Rothschild, Paris, France; Department of Neurosurgery, University Hospital Bonn, Bonn, Germany

## Abstract

Photo-activatable drugs (PDs) are rapidly emerging as precision therapeutics across biomedical research, yet their potential in epilepsy treatment has remained understudied. Given that 30% of epilepsies are medically refractory, and anti-seizure medications often cause multi-organ side-effects, PDs could break new ground. Here, we evaluate light-switchable ion-channel blockers QAQ and CQAQ, and a newly developed caged propofol (CaP), in murine brain slices, and postsurgical brain tissue from patients with epilepsy or brain tumors. In mice, we show that QAQ/CQAQ reversibly suppress neuronal firing, while photo-activated CaP prolonged inhibitory post-synaptic currents and enhanced leak current. While QAQ caused identical effects in human tissue, CQAQ unexpectedly increased firing. Activated CaP robustly suppressed epileptiform activity across species. This work establishes CaP as a tool for neuroscience and disease-related studies. Our results highlight the necessity for early human model testing in biomedical research, and showcase photopharmacology as a potential powerful approach to control human epilepsy.

## Introduction

Beyond phototherapy and photodynamic therapy long applied in the clinical context(*1*, *2*), photo-activatable drugs (PDs) are rapidly emerging as precision therapeutics across diverse biomedical fields(*3*), including treatment of various forms of cancer(*4*), cardiac arrhythmia(*5*), pain disorders(*6*, *7*), or for vision restoration(*8*, *9*). For the latter, PDs are at the brink of clinical transfer(*10*). Despite this promise, few studies have explored PDs in epilepsy research(*11–14*), and none have tested them in experimental human epilepsy models. Epilepsy, a common and often debilitating neurological disorder based on aberrant neuronal networks affecting around 1% of the general population(*15*), remains uncontrolled in up to one third of patients, despite a growing arsenal of anti-seizure medications (ASM). This so-called refractory epilepsy poses an immense burden to those suffering from it, as the risk of seizure occurrence prevents patients from participating in basic essential activities of daily life, like swimming, or driving a car. Moreover, more than half of patients experience significant pharmacotherapy-related side effects, most notably psychiatric, cardiac, hepatotoxic, gastro-intestinal or skin-related symptoms(*16*).

With regards to both treatment efficacy and tolerability in refractory focal epilepsy, PDs could break new ground. PDs can be locally activated on demand in single or multifocal epileptic brain areas – including those that are surgically inaccessible or functionally indispensable, and therefore unsuitable for resection. The spatially confined activation minimizes systemic exposure and side effects, potentially enabling the re-introduction of potent compounds that were previously withdrawn from clinical use due to unfavorable systemic side-effects (*13*, *17*). Further, the PD strategy allows the adoption of widely used potent drugs with known anti-seizure capacity currently used for other indications. For example, general anesthetics – commonly administered to treat status epilepticus in intensive care – could be locally activated with a PD strategy, extending their utility to epileptology outpatient settings. Importantly, different PDs can be flexibly combined or exchanged, and the approach does not require gene transfer(*3*).

Here, we use light-switchable ion-channel blockers QAQ and CQAQ(*6*, *18*), and a newly developed caged propofol (CaP) in acute mouse brain slices, and post-surgical human brain tissue from patients with refractory focal epilepsy or brain tumors. Using patch-clamp and field potential recordings in murine slices, we show that QAQ and the dark-inactive CQAQ reversibly suppress neuronal firing in a wavelength-dependent manner. CaP markedly prolongs inhibitory post-synaptic potentials and increases leak current, also in a wavelength-dependent fashion. Intriguingly, in human brain tissue, while QAQ showed consistent effects, CQAQ unexpectedly increased firing rate upon light-activation. We further evaluated CaP in murine and human brain slices under epileptiform conditions, and show that it efficiently blocks interictal epileptiform activity (IEA), and seizure-like episodes (SLE). Our results showcase photopharmacology as a versatile tool to control refractory focal epilepsy. This approach could reduce the need for surgical resection, and may be extended to multi-focal epilepsy using minimally invasive, biocompatible light-fiber implants(*19*).

## Results

### Potent anesthetics as photo-activatable anti-seizure drugs

Our goal was to study the potential of switchable and caged PDs in human epilepsy, with a focus on anesthetics. Lidocaine is an ion channel blocker mostly applied for local anesthesia of peripheral nerves. In a dose-dependent fashion, its systemic application can encompass potentially serious cardiotoxic and neurological side effects in humans(*20*), and has been shown to trigger lethal side effects in rodent models at higher doses(*21*). Light-activatable lidocaine could overcome these issues, and indeed photo-switchable QAQ and sign-inverted CQAQ have been previously developed, but not tested in epilepsy research. Further, we sought to develop a light-activatable propofol, the most-widely used general anesthetic in medicine(*22*). In operative and intensive care, its effect of loss of consciousness upon systemic injection is a desired action, and it is also used as the last line in treating refractory status epilepticus. Yet, due to its action profile and undesired side-effects(*22*), propofol is currently not applicable as an anti-seizure medication in the outpatient setting.

### Development and characterization of caged propofol

We set out to develop light-activatable propofol as a caged compound. To this end, we aimed to couple a *m*,*p*-dimethoxy-*o*-nitrobenzyl residue as a photocleavable protecting group (PPG) to the phenolic function of propofol (Fig. 1 A). Initially, we prepared a direct ether linkage (Caged Propofol version 1, ‘CaP-v1’), or connected propofol via a carbonate ester to the PPG (CaP-v2, Fig. 1 A). However, both conjugates were too stable resulting in only low cleavage rates upon irradiation at 365 or 400 nm wavelength, respectively (Fig. 1 B). Thus, we introduced an acetal linker between propofol’s phenolic group and the PPG similar as in the water-soluble prodrug fospropofol(*23*) yielding CaP-v3, which was further optimized by methyl substitution to obtain a branched PPG derivative, and by replacing the methoxy groups by a methylenedioxy substitution (CaP-v4, Fig. 1 A). This led to high cleavage rates upon irradiation (Fig. 1 B), and CaP-v4 was thus suitable for pharmacological studies. As of here, we refer to CaP-v4 as CaP (for synthesis details of CaP-v1 to -v4, see methods). As expected, cleavage of CaP releasing propofol was dependent on the wavelength of irradiation, with a suitable range from 365 to 420 nm (Fig. 1 C, D). Besides CaP, we further evaluated two previously reported photoswitchable lidocaine derivatives, quaternary ammonium-azobenzene-quaternary ammonium (QAQ) connected by amide bonds, and CQAQ, a cyclic diazocine version of QAQ (Fig. 1 E)(*6*, *18*). Similarly to switchable propofol reported previously(*24*), QAQ is *trans*-configurated (“active”) in the dark undergoing a UV light-induced switch to its inactive *cis*-stereoisomer, while CQAQ displays *cis*-configuration in its dark-adapted state, and is switched to the extended *trans* form by UV light. In turn, green light activates QAQ, while it inactivates CQAQ. Thus, both QAQ and CQAQ allow fast and reversible ion channel blockade upon wavelength-specific irradiation in the green and UV range of light.

**Figure 1:**
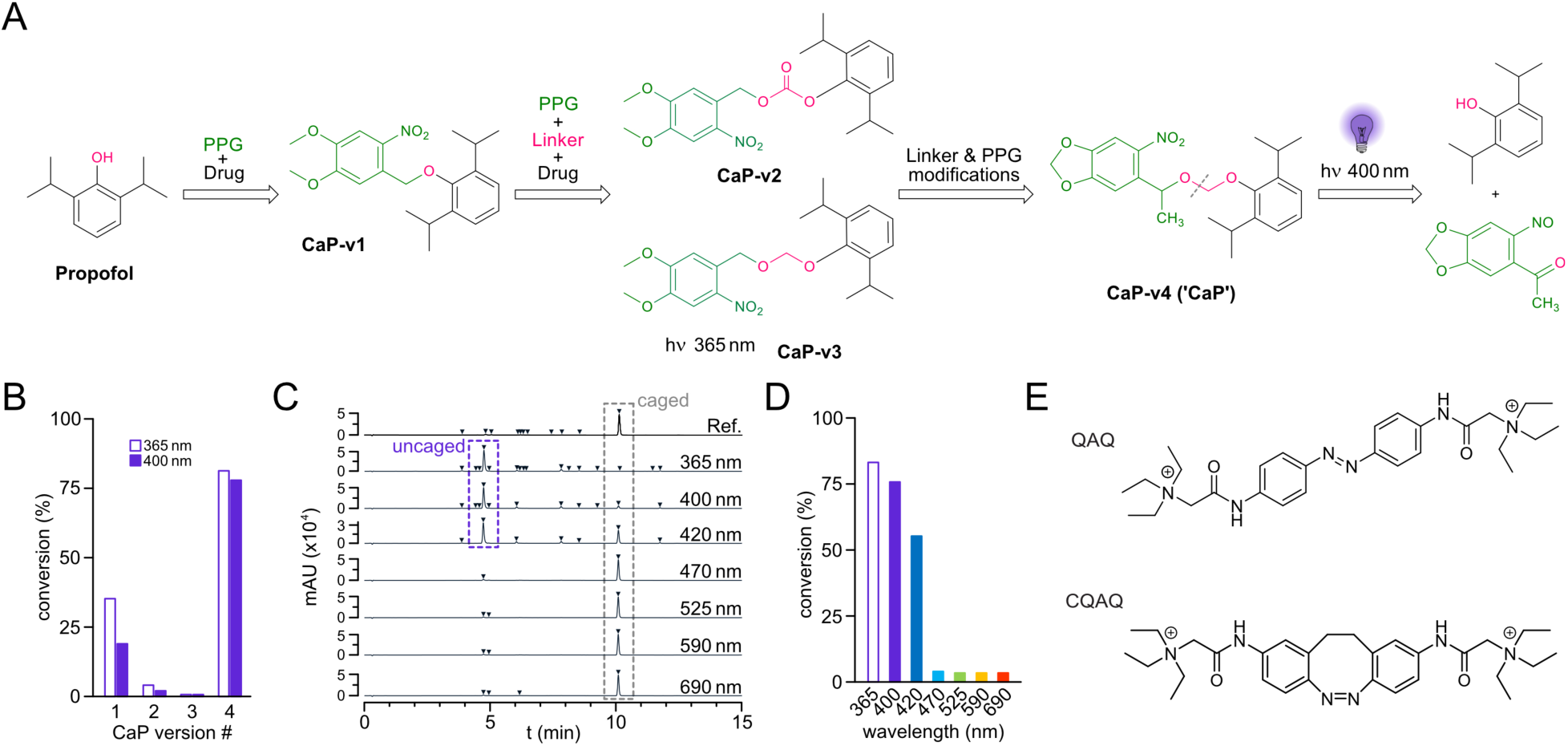
Synthesis and characterization of caged propofol, QAQ and CQAQ. **A)** Synthesis of caged propofol versions 1 to 4 (CaP-v1/-v2/-v3/-v4). At CaP-v4 (‘CaP’), the dotted gray line indicates where the bond is broken by irradiation. **B)** LCMS-measured conversion of different caged propofol versions upon wavelength-specific irradiation (365 or 400 nm). **C)** LCMS-measured conversion of CaP-v4 (‘CaP’) upon irradiation across a wide range of wavelengths. Note efficient conversion between 365 to 420 nm. mAU: milli-absorbance unit **D)** Same as C, displaying conversion efficiency of CaP across different wavelengths. **E)** Structures of QAQ and CQAQ.

### Basic functional evaluation of QAQ, CQAQ and CaP in brain tissue

To evaluate QAQ, CQAQ and the newly developed CaP action on brain tissue, we first carried out neuronal patch-clamp recordings in mouse brain slices following two experimental protocols (Fig. 2 A, Suppl. Fig. 1 A). For QAQ and CQAQ that have to be applied intracellularly (except under certain conditions where they may enter cells via P2X-receptors(*25*)), we performed whole-cell patch clamp recordings of individual hippocampal neurons (CA1 region, see methods for details) in submerged brain slices with the PDs being dissolved in the internal patch solution. In this setting, we carried out repeated 1-second sweeps of direct current injection (120-700 pA) to trigger neuronal action potential firing in the presence of different wavelengths of light that either activate or inactivate QAQ or CQAQ. For extracellularly applied CaP, which exerts its action mainly via a modulation of synaptic transmission, we carried out stimulation (0.2 ms, 100-530 µA) of the Schaffer collaterals (input fibers to the CA1 region) using an additional bipolar electrode, while recording inhibitory post-synaptic currents (IPSCs) and leak current in postsynaptic CA1 neurons (see methods for details).

**Figure 2:**
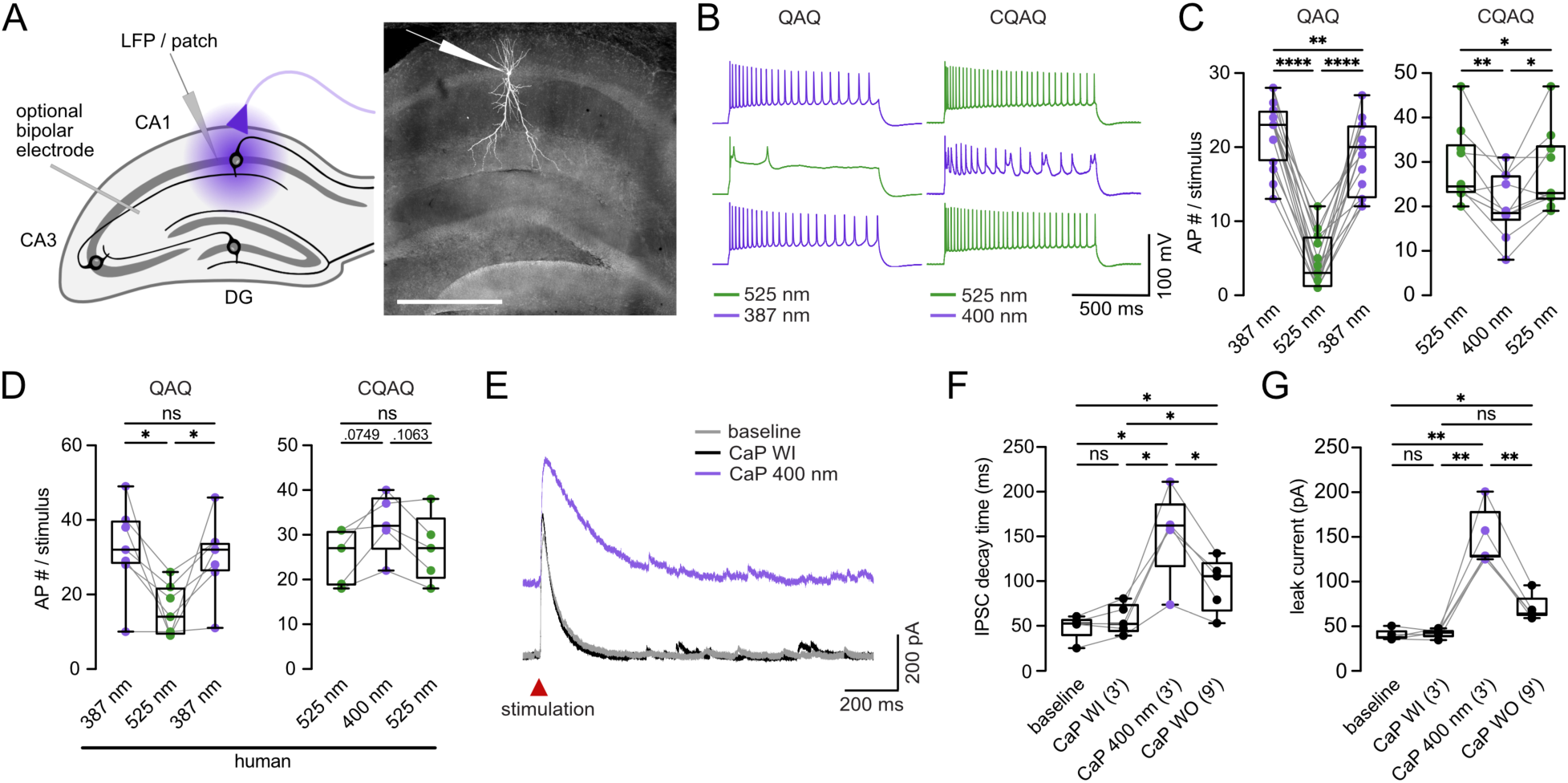
Light-dependent effects of QAQ, CQAQ and caged propofol (CaP) on neuronal activity in mouse and human brain slices. **A)** Left: Scheme of experimental setup for patch-clamp and local field potential (LFP) recordings in coronal mouse hippocampal slices. Right: Biocytin staining of a CA1 neuron in a coronal mouse hippocampal slice co-stained with DAPI. Typical recording location marked by pipette. Scale bar: 500 µm. DG: dentate gyrus. **B)** Representative current clamp recordings showing the light-dependent effect of QAQ (left) and CQAQ (right) on action potential (AP) firing rate in mouse hippocampal neurons (CA1). Green and violet colors represent illumination wavelengths: 525 nm (QAQ activation, CQAQ inactivation), 387 nm (QAQ inactivation), 400 nm (CQAQ activation). **C)** Quantification of the effect and effect-reversibility of QAQ and CQAQ on stimulus-evoked (1 sec, 120 - 700 pA, applied through patch pipette) AP firing rate in mouse hippocampal neurons (CA1). Left: QAQ (15 cells, 11 mice), 22.00 ± 1.15 (387 nm), 4.40 ± 0.92 (525 nm), 18.80 ± 1.26 (2^nd^ 387 nm). Repeated measures (RM) one-way ANOVA + Tukey’s test (F [1.332, 18.647] = 127.352): 387 nm vs. 525 nm p < 0.0001, 525 nm vs. 2^nd^ 387 nm p < 0.0001, 387 nm vs. 2^nd^ 387 nm p = 0.0039. Right: CQAQ (10 cells, 6 mice), 28.70 ± 2.67 (525 nm), 20.40 ± 2.23 (400 nm), 27.60 ± 2.84 (2^nd^ 525 nm). RM one-way ANOVA + Tukey’s test (F [1.045, 9.409] = 13.245): 525 nm vs. 400 nm p = 0.0081, 400 nm vs. 2^nd^ 525 nm p = 0.0220, 525 nm vs. 2^nd^ 525 nm p = 0.0419). **D)** Quantification of the effect and effect-reversibility of QAQ and CQAQ on stimulus evoked (1 sec, 150 - 750 pA, applied through patch pipette) AP firing rate in human hippocampal neurons (dentate gyrus [DG]). Left: QAQ (7 cells, 5 patients), 32.29 ± 4.62 (378 nm), 15.57 ± 2.59 (525 nm), 30.14 ± 4.00 (2^nd^ 378nm). RM one-way ANOVA + Tukey’s test (F [1.073, 6.438] = 10.443): 387 nm vs. 525 nm p = 0.0384, 525 nm vs. 2^nd^ 387 nm p = 0.0402, 387 nm vs. 2^nd^ 387 nm p = 0.2630. Right: CQAQ (5 cells, 2 patients), 25.20 ± 2.84 (525 nm), 32.40 ± 3.08 (400nm), 27.00 ± 3.44 (2^nd^ 525nm). RM one-way ANOVA + Tukey’s Test (F [1.848, 7.393] = 6.718): 525 nm vs. 400 nm p = 0.0749, 400 nm vs. 2^nd^ 525 nm p = 0.1063, 525 nm vs. 2^nd^ 525 nm p = 0.6232). Note opposite effect of CQAQ on human vs. mouse hippocampal neurons. **E)** Representative voltage clamp recordings showing the light-dependent effect of CaP on inhibitory postsynaptic current (IPSC) decay time and leak current in mouse hippocampal neurons (CA1). Note shift in baseline upon CaP activation reflecting change in leak current. WI: wash in. **F-G)** Quantification of the light-dependent effect and effect-reversibility of CaP on stimulus-evoked (0.2 ms, 290 - 500 µA, applied via bipolar electrode in Schaffer collaterals) IPSC decay time and leak-current. **F)** IPSC decay time (ms) in mouse hippocampal neurons (CA1; 5 cells, 5 mice): 48.99 ± 6.18 (baseline), 57.46 ± 7.51 (CaP WI), 153.5 ± 22.21 (CaP 400nm), 95.99 ± 13.61 (CaP WO). RM one-way ANOVA + Tukey’s test (F [1.249, 4.995] = 28.484): baseline vs. CaP WI p = 0.3914, CaP WI vs. CaP 400 nm p = 0.0225, CaP 400 nm vs. CaP WO p = 0.0252, baseline vs. CaP 400 nm p = 0.0127, baseline vs. CaP WO p = 0.0226, CaP WI vs. CaP WO p = 0.0383. WI: wash in, WO: wash out. **G)** Leak current (pA) in mouse hippocampal neurons (CA1; 5 cells, 5 mice): 40.24 ± 2.74 (baseline), 42.09 ± 2.17 (CaP WI), 148.15 ± 14.36 (CaP 400 nm), 70.34 ± 6.55 (CaP WO). RM one-way ANOVA + Tukey’s test (F [1.137, 4.548] = 51.346): baseline vs. CaP WI p = 0.9173, CaP WI vs. CaP 400 nm p = 0.0060, CaP 400 nm vs. CaP WO p = 0.0025, baseline vs. CaP 400 nm p = 0.0062, baseline vs. CaP WO p = 0.0489, CaP WI vs. CaP WO p = 0.0523. For entire fig.: Boxes extend from 25^th^ – 75^th^ percentiles, inner line represents the median. All ± represent s.e.m. Depiction of statistical significance: n.s. not significant, *p < 0.05, **p < 0.01, ***p < 0.001, ****p < 0.0001.

Confirming previous reports(*6*, *18*), both QAQ and CQAQ allowed reversible blockade of action potential firing in a wavelength-dependent fashion in mouse brain slices (Fig. 2 B, C). In post-resective brain tissue from patients with refractory epilepsy, and patients with brain tumors without clinical epilepsy diagnosis (for cohort details, see suppl. table 1), QAQ again led to a profound and reversible decrease in neuronal firing rate (Fig. 2 D, left). Unexpectedly, opposite to our murine recordings, CQAQ reversibly increased firing upon light activation (Fig. 2 D, right). Both in murine and human tissue, light stimulation alone did not change AP rate nor input resistance upon stimulation (Suppl. Fig. 1 B). In addition to these results, we confirmed that QAQ and CQAQ have differential characteristics, in keeping with previous descriptions (Suppl. Fig. 2 A-E)(*18*). Unlike QAQ that reversed back into its preferred active *trans*-state within just a few minutes in the dark, CQAQ slowly inactivated in the dark making it a more suitable candidate for in vivo use in this regard (Suppl. Fig. 2 E, F). In turn, QAQ had a significantly stronger effect size on action potential suppression than CQAQ (Suppl. Fig. 2 G), likely in part due to the different switching properties of the two molecules at similar light intensities (Suppl. Table 2).

After initial evaluation of QAQ and CQAQ, we moved on to assess functionality of our newly developed CaP molecule in brain tissue. In line with the published mechanism of propofol, CaP led to a profound shift of leak current and an increase in IPSC decay time (Fig. 2 E-G) upon wavelength-dependent uncaging at biocompatible light-intensities (153.18 mW/cm^2^), with a similar effect size to native propofol application (Suppl. Fig. 3 A). Light application in the presence of the photoprotective group alone did not change leak current or IPSC decay time (Suppl. Fig. 3 B). Notably, the light-dependent CaP effects partially recovered upon extended washout (Fig. 2 F, G). Together, these experiments showed that both switchable and caged anesthetics can be used for efficient control of neuronal activity across species (mouse, human), both in the healthy brain and in chronic neurological diseases (epilepsy, brain tumors).

### Suppression of epileptiform activity in mouse and human brain tissue

After demonstrating efficacy in control of neuronal activity using QAQ, CQAQ and CaP, we next tested if CaP activation can block epileptiform activity in mouse and human brain tissue at the network level. In principle, both CQAQ and CaP would technically be amenable to in vivo application, as both are inactive in the dark. Yet, based on the intracellular target mechanism of CQAQ, its switching efficiency (Suppl. Table 2), and its unexpected effect on neuronal firing in human brain tissue, we decided to focus on CaP for the following experiments. In an interface chamber setting, we first tested CaP in murine brain slices under acutely induced epileptiform activity (low Mg^2+^ model, see methods for details). We performed continuous local field potential (LFP) recordings across the establishment of steady-state epileptiform activity, wash-in of CaP without light-activation (4 min), light activation (6 min), and wash-out. Similar to the native propofol positive controls without activation of light, both interictal epileptiform activity (IEA, Fig. 3 A, C) and seizure-like episodes (SLE, Fig. 3 B, D) could be blocked by CaP in a light-dependent fashion. Control experiments were performed using either CaP without light-activation, or light application alone, and did not yield any reduction of epileptiform activity (Fig. 3 C-D, Suppl. Fig. 3 C). Likewise, light application in the presence of the photoprotective group alone did not block IEA or SLE (Suppl. Fig. 3 D). As in the experiments under physiological conditions, the light-dependent CaP effects recovered upon extended washout (Suppl. Fig. 3 C, top panel).

**Figure 3:**
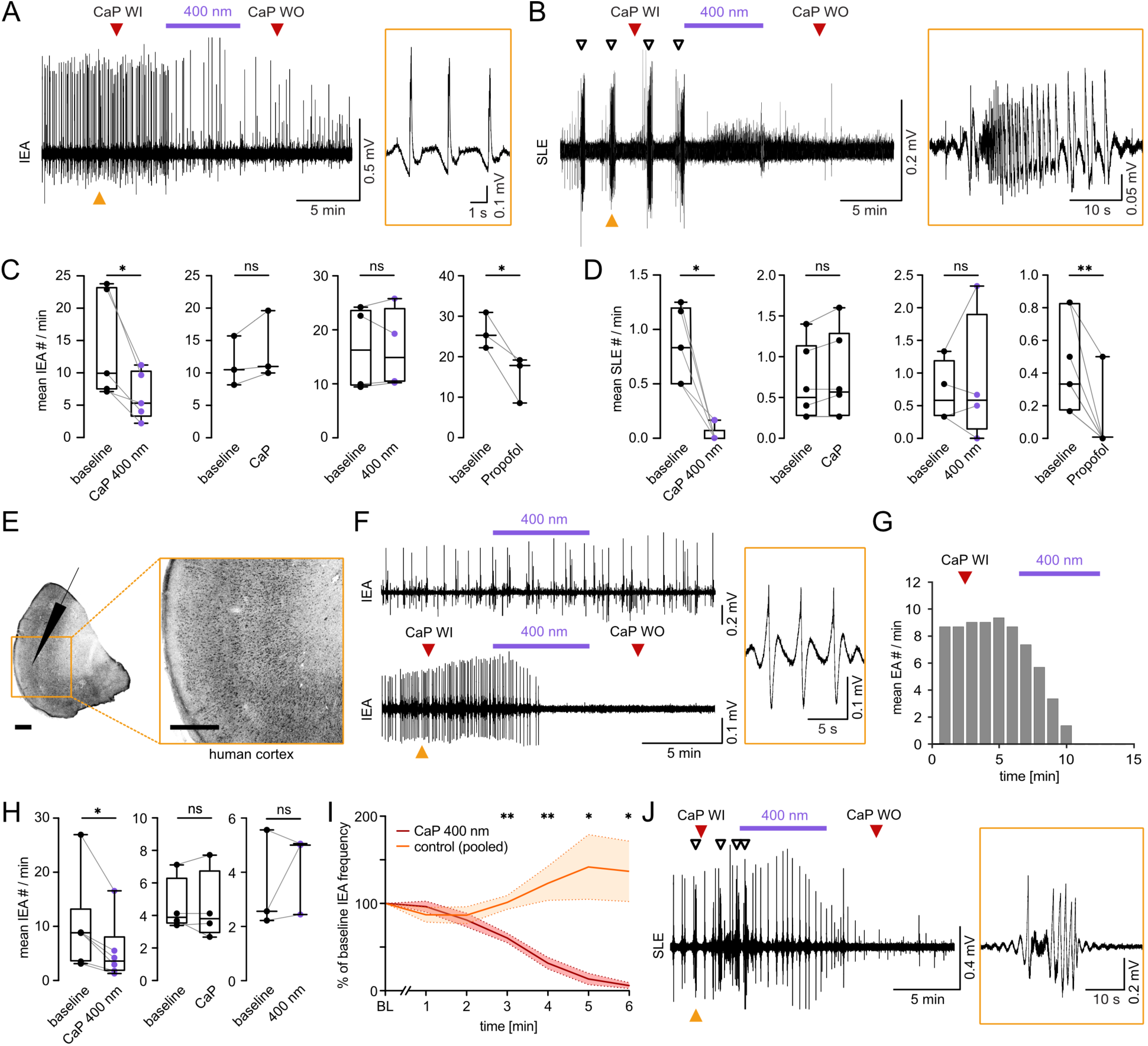
Light-activated CaP inhibits epileptiform activity in mouse and human brain slices. Interictal epileptiform activity (IEA) and seizure-like episodes (SLE) were evoked by Mg^2+^-free ACSF solution in both murine (A-D) and human (E-J) cortical slices. **A)** Left: Representative LFP recording showing the light-dependent effect of CaP on event number (#) of IEA. WI: wash in; WO: wash out. Violet color represents illumination at 400 nm (CaP activation). Yellow inset: zoom-in of three IEA events (at yellow arrow in LFP trace). **B)** Representative LFP recording showing the light-dependent effect of CaP on # of SLE (black arrows). Yellow inset: zoom-in of one SLE (at yellow arrow in LFP trace). **C)** Quantification of the effect of light-activated CaP, CaP without light-activation or light-only application (neg. controls), and native propofol (pos. control) on IEA #. Left (5 slices, 4 mice): 14.27 ± 3.75 (baseline), 6.50 ± 1.70 (CaP 400 nm). Paired t-test: df = 4; p = 0.0260. Center left (3 slices, 3 mice): 11.46 ± 2.23 (baseline), 13.54 ± 3.05 (CaP [no light]). Paired t-test: df = 2; p = 0.1702. Center right (4 slices, 2 mice): 16.55 ± 3.98 (baseline), 16.45 ± 3.76 (400 nm [light-only]). Paired t-test: df = 3; p = 0.9280. Right (3 slices, 2 mice): 26.15 ± 2.55 (baseline), 15.19 ± 3.34 (native propofol). Paired t-test: df = 2; p = 0.0460. **D)** Quantification of the effect of light-activated CaP, CaP without light-activation or light-only application (neg. controls), and native propofol (pos. control) on SLE #. Left (5 slices, 3 mice): 0.85 ± 0.16 (baseline), 0.03 ± 0.03 (CaP 400 nm). Paired t-test: df = 4; p = 0.0104. Center left (6 slices, 4 mice): 0.83 ± 0.24 (baseline), 0.93 ± 0.28 (CaP [no light]). Paired t-test: df = 5; p = 0.0842. Center right (4 slices, 3 mice): 0.71 ± 0.24 (baseline), 0.88 ± 0.51 (400 nm, [light-only]). Paired t-test: df = 3; p = 0.6134. Right (7 slices, 4 mice): 0.45 ± 0.11 (baseline), 0.07 ± 0.07 (native propofol). Paired t-test: df = 6; p = 0.0050. **E)** NeuN staining of a human cortical slice. Typical recording location marked by pipette. Yellow inset: Magnification of recording location. Scale bar: 1 mm. **F)** Representative LFP recordings in human cortical tissue; top: control experiment, light application alone does not reduce IEA; bottom: light-dependent effect of CaP on IEA #. Yellow inset: zoom-in of three IEA events (at yellow arrow in LFP trace). **G)** Same experiment as F: histogram (bin size: 1 min, running average over three bins) showing the time-dependent effect of ligh-activated CaP on IEA #. **H)** Quantification of the effect of light-activated CaP, CaP without light-activation or light-only application on IEA #. Left (6 slices, 4 patients): 10.01 ± 5.38 (baseline), 5.38 ± 2.32 (CaP 400 nm). Paired t-test: df = 5, p = 0.0175. Center (4 slices, 3 patients): 4.57 ± 0.86 (baseline), 4.50 ± 1.11 (CaP [no light]). Paired t-test: df = 3, p = 0.8428. Right (3 slices, 2 patients): 3.45 ± 1.06 (baseline), 4.17 ± 0.86 (400 nm [light only]). Paired t-test: df = 2, p = 0.5012. **I)** Quantification of IEA frequency across 6 min of light-activated CaP (6 slices, 4 patients) vs. pooled CaP (no light; 4 slices, 3 patients) and 400 nm (light only; 3 slices, 2 patients) controls. Multiple unpaired t-tests + Holm-Šídák correction: min1: 96.34 ± 6.89 (CaP) vs. 87.07 ± 8.46 (control), p = 0.6679; min2: 80.96 ± 8.69 (CaP) vs. 86.58 ± 9.62 (control), p = 0.6774; min3: 60.36 ± 5.95 (CaP) vs. 101.27 ± 7.86 (control), p = 0.0098; min4: 31.53 ± 7.27 (CaP) vs. 123.01 ± 19.48 (control), p = 0.0080; min5: 13.41 ± 7.12 (CaP) vs. 141.76 ± 37.08 (control), p = 0.0278; min6: 5.80 ± 3.67 (CaP) vs. 136.78 ± 34.56 (control), p = 0.0207. Dotted lines/shades represent s.e.m. values. Note IEA suppression upon CaP light-activation compared to increase of IEA # across time in control group. BL: baseline. **J)** LFP recording showing the light-dependent effect of CaP on # of SLE (black arrows) in human cortical tissue. Yellow inset: zoom-in of one SLE (at yellow arrow in LFP trace). Note rapid cessation of SLE upon CaP light-activation, and subsequent complete suppression of IEA. For entire fig.: Boxes extend from 25^th^ – 75^th^ percentiles, inner line represents the median. All ± represent s.e.m. Depiction of statistical significance: n.s. not significant, *p < 0.05, **p < 0.01, ***p < 0.001, ****p < 0.0001.

Finally, we tested our CaP approach under epileptic conditions (low Mg^2+^ model) in acute post-resective human brain tissue from patients with refractory focal epilepsy or brain tumors (Fig. 3 E, for cohort details, see Suppl. Table 3). As in the murine recordings, native propofol application led to a reduction of IEA and SLE (Suppl. Fig. 3 E-F). Similarly, light-activated CaP led to a massive reduction of IEA, while no such effect was found in controls (Fig. 3 F-I, Suppl. Fig. 3 G). IEA suppression increased over the course of minutes, with near complete cessation of pathological network events at around 5 min after the start of photo-activation (Fig. 3 F, G, I). In one case of refractory human epilepsy, we also succeeded in recording SLE in post-resective brain tissue (Fig. 3 J). Here too, CaP activation led to a complete cessation of SLE (Fig. 3 J, Suppl Fig. 3 H). In sum, these experiments proved that CaP efficiently blocks epileptiform activity in a light-dependent fashion in mouse brain slices, and post-resective human brain tissue from patients with refractory epilepsy.

## Discussion

In this study, we show that neuronal activity in chronic epilepsy can be efficiently suppressed using biocompatible light-intensities and biotolerable irradiation periods (up to a few minutes). This was achieved using two previously established photo-switchable ion channel blockers (QAQ, CQAQ), as well as a newly developed caged propofol (CaP). Importantly, we provide direct evidence in human brain tissue that targeted photopharmacology could serve as a precise and effective strategy for human refractory epilepsy. A photopharmacological approach to focal epilepsy offers several key advantages. First, compared to resective surgery, it is minimally invasive, requiring merely the implantation of biocompatible light-fibers to the target region(*19*). This minimal invasiveness allows application of targeted photopharmacology in cases where surgery is not an option, as in certain multi-focal epilepsies, involvement of eloquent brain regions, or surgically inaccessible brain regions. Similar to recent optogenetic approaches, photopharmacology could be employed using closed-loop intervention(*26*, *27*). Yet, beyond non-reliance on viral gene transfer, photopharmacology also offers flexible use of PDs with different mechanisms of action, which can be changed or combined using the same light source.

This study has inherent limitations. All compounds used here require light-stimulation in the UV to low blue range, which would have to be redesigned for activation at longer wavelength to enable clinical transfer. Red-shifting the photoactivation would be necessary both to improve biotolerability and light penetration depth in biological tissue. While these efforts are ongoing, optogenetic studies in mice have shown that sufficiently red-shifted compounds (excitation >600 nm) allow targeted stimulation of deep brain regions (depths up to 7 mm) through the intact skull(*28*), and several ex-vivo studies in human brain tissue have demonstrated that red/near-infrared light can reach brain depths up to the centimeter range(*29*, *30*). These findings suggest that, in humans, clinically relevant brain volumes – including focal seizure-generating networks – could be reached via multiplexed light outlets distributed along or around implanted optical probes.

Moreover, the photocleavable prodrug CaP developed in this study remains an experimental drug. For clinical application, it would have to undergo extensive optimization, not only to achieve activation at longer wavelengths, but also to refine its physicochemical, pharmacokinetic and toxicological properties. In vitro, photo-activated CaP reached its maximum effect size on epileptiform network activity over minutes. This would be fast enough to disrupt seizure clusters or status epilepticus, but it remains to be determined whether initial effects sizes of photo-activated CaP suffice for rapid on demand seizure control. Finally, in the in-vitro experiments presented here, all photoactivatable compounds were delivered locally. For use in intact organisms, peroral or systemic routes of administration would be desirable, which again will require further optimization. The small size of propofol makes it a particularly appealing candidate for the photo-prodrug approach pursued in this study.

Our primary goal regarding CaP was to develop a potent light-activatable drug to tackle hard-to-treat epilepsy. Beyond this specific goal, however, our work has broader implications for both basic neuroscience and other medical fields. Because the CaP uncaging wavelength lies between ∼365-420 nm, it could be used in basic research that involves simultaneous functional fluorescence imaging, with sufficient spectral separation(*31–35*). This also ties in with other biomedical disciplines such as cancer neuroscience. It has been well established over recent years that astrocytic brain tumors form a multi-cellular functional network, and that the activity of neurons forming part of this network drives tumor growth(*36*, *37*). Here, photoactivatable anesthetics such as CaP could be useful in locally disrupting tumor cell network communication, potentially curbing tumor progression. Finally, given that chronic depth electrode implants with closed-loop stimulation capabilities have been used in several neurological disorders including epilepsy(*38*, *39*), their integration with on demand biocompatible light delivery systems is not farfetched(*19*, *40–43*). Combined with the fact that targeted photopharmacology does not rely on gene transfer, this approach could be clinically tested in the near future, representing a realistic pathway toward minimally invasive, precision therapy for refractory epilepsy.

## Methods

### Experimental models and subject details

#### Animals and animal protocols

The study design as well as the reporting of results in this paper were based on the ARRIVE guidelines and 3R principles. All procedures were carried out with care and in compliance with the European, German national, and institutional regulations (animal protocol 81-02.04.2024.A009). Animals were housed at a 12 h inverted light cycle (12 hours of darkness from 7 am – 7 pm, 12 hours of light), and food and water were accessible ad libitum. Adult male and female wild type C57BL/6J at the age of 8-14 weeks were used for experiments.

#### Patient data

Hippocampal or cortical tissue blocks were obtained directly following neurosurgical resection in 17 patients (epileptological and/or neuro-oncological cases, age 19 – 77 years, for details see suppl. tables 1 and 3), at the Dept. of Neurosurgery at University Medical Center Bonn. Informed compliance was signed by all patients, and ethical approval (# 078/22) for the study obtained from the Medical Institutional Review Board of the University of Bonn.

### Synthesis and characterization of photoactivatable compounds

#### Synthetic procedures

Caged propofol, version 1 (CaP-v1). 1-(((2,6-Diisopropylphenoxy)methoxy)methyl)-4,5-dimethoxy-2-nitrobenzene (**CaP-v1**) was obtained by alkylation of propofol with 1-(bromomethyl)-4,5-dimethoxy-2-nitrobenzene (**1**) in the presence of potassium carbonate in dry dimethylformamide (DMF).

**CaP-v1** underwent photolysis to release propofol with moderate efficiency (35% at 365 nm, 19% at 400 nm within 2 min), demonstrating proof-of-concept for light-triggered release while highlighting the need for further structural optimization to improve conversion under short irradiation times (Fig. 1B).

**Scheme 1.**
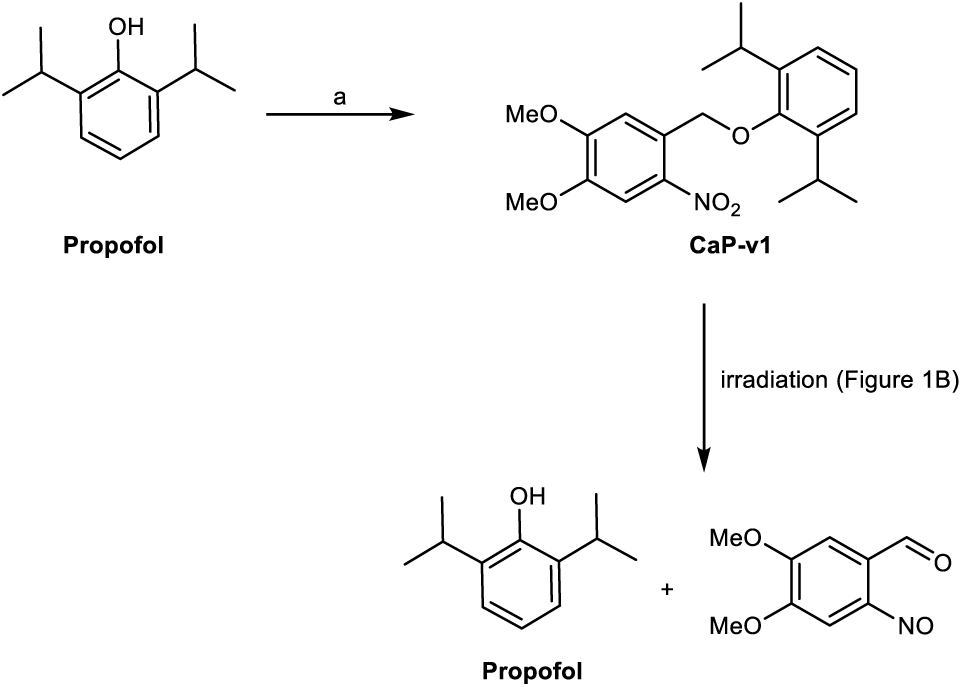
Synthesis of 1-((2,6-diisopropylphenoxy)methyl)-4,5-dimethoxy-2-nitrobenzene **(CaP-v1)**. a. 1-(Bromomethyl)-4,5-dimethoxy-2-nitrobenzene (1), K_2_CO_3_, DMF, RT, 14 h.

Caged propofol, version 2 (CaP-v2). 2,6-Diisopropylphenyl (4,5-dimethoxy-2-nitrobenzyl) carbonate (**CaP-v2**) was prepared in two steps. (4,5-dimethoxy-2-nitrophenyl)methanol (**2**) was converted to the corresponding carbonochloridate **3** via reaction with triphosgene in the presence of triethylamine(*44*). Then, propofol was coupled with **3**. **CaP-v2**, featuring a carbonate linker between propofol and the photocleavable protecting group, exhibited a very low degree of photocleavage (4% at 365 nm and 2% at 400 nm within 2 min), indicating that the carbonate linkage conferred increased photostability compared to **CaP-v1**. It was thus not suitable for achieving rapid, high-yielding photolysis with short irradiation times (Fig. 1B).

**Scheme 2.**
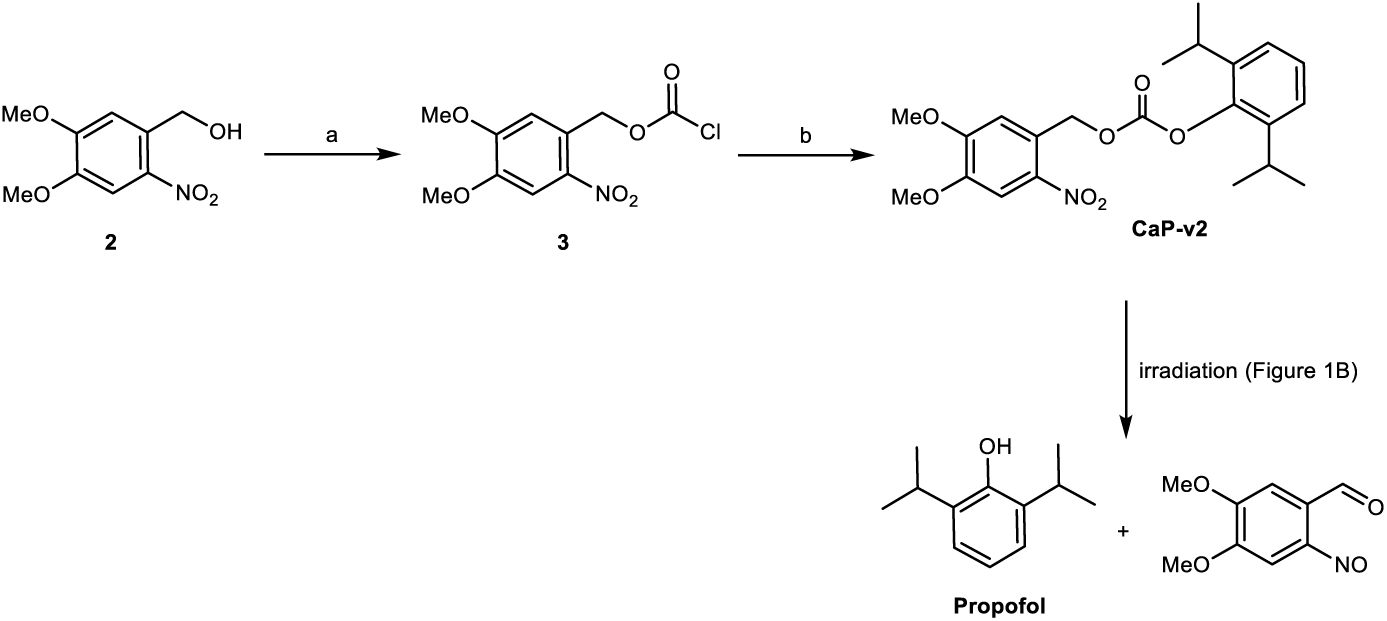
Synthesis of 2,6-diisopropylphenyl (4,5-dimethoxy-2-nitrobenzyl) carbonate **(CaP-v2)**. a. Triphosgene, Et_3_N, THF, 18h. b. Propofol, Et_3_N, DCM, 2.5 h, 0 °C to RT

Caged propofol, version 3 (CaP-v3). 1-(((2,6-Diisopropylphenoxy)methoxy)methyl)-4,5-dimethoxy-2-nitrobenzene (**CaP-v3**) was synthesized in three steps. In the first step, propofol was alkylated with chloromethyl methyl sulfide under basic conditions in hexamethylphosphoramide (HMPA) to afford the methylsulfane **4**. Oxidative chlorination of **4** with thionyl chloride produced the benzyl chloride **5** (*45*). Then, 4,5-dimethoxy-2-nitrophenyl)methanol (**2**) was deprotonated with sodium hydride in HMPA and coupled with **5** to furnish **CaP-v3**. This step proved challenging requiring extensive purification involving both flash chromatography and preparative TLC. Structurally, **CaP-v3** incorporates a methylene linker between propofol and the photocleavable protecting group. As shown in figure 1B, this compound was resistant to photolysis, with no detectable cleavage under the tested irradiation conditions.

**Scheme 3.**
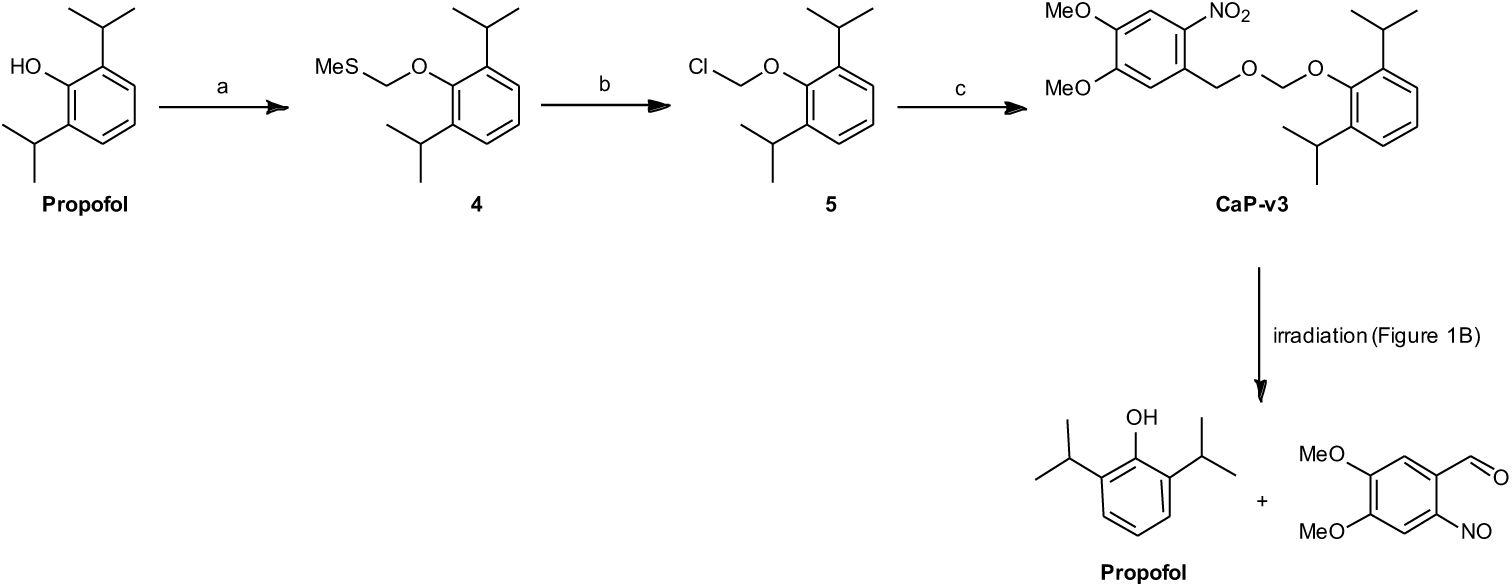
Synthesis of 1-(((2,6-diisopropylphenoxy)methoxy)methyl)-4,5-dimethoxy-2-nitrobenzene **(CaP-v3)**. a. Chloromethyl methyl sulfide, NaH, HMPA, RT, 18h. b. SO₂Cl₂, DCM, 0 °C to RT, 3h. c. (4,5-Dimethoxy-2-nitrophenyl)methanol (**2**), NaH, HMPA, RT, 18h.

Caged propofol, version 4 (CaP-v4, also termed ‘CaP’). (*R,S*)-5-(1-((2,6-diisopropylphenoxy)methoxy)ethyl)-6-nitrobenzo[*d*][1,3]dioxole (**CaP-v4**) was synthesized in five steps. The synthesis of 5-(1-(chloromethoxy)ethyl)-6-nitrobenzo[*d*][1,3]dioxole (**11**) was adapted from a previously reported procedure(*46*) with major modifications. Benzodioxolylethenone **6** was treated with nitric acid (HNO_3_) in nitromethane to yield nitrobenzodioxolylethenone **7**. Reduction of **7** with sodium borohydride provided the corresponding nitrobenzodioxolylethanol **8**. Alkylation of **8** afforded the ether **10**, which was subsequently converted to the corresponding chloride **11** by treatment with sulfuryl chloride. In the final step, propofol was alkylated with **11** under basic conditions to afford **CaP-v4**.

Structurally, the photocleavable protecting group in **CaP-v4** contains a methyl substituent at the benzylic position and a dioxole ring annelated to the nitrobenzyl residue. As shown in figure 1B, this led to the highest rate of photocleavage in the present series, with conversions of 81% at 365 nm and 78% at 400 nm within 2 min. The narrow and efficient activation profile between 365 nm and 420 nm, combined with minimal background release at longer wavelengths, establishes **CaP-v4** as the lead candidate for light-controlled propofol delivery.

1-(6-Nitrosobenzo[*d*][1,3]dioxol-5-yl)ethan-1-one (**9**). The photocleavage product **9**, required as a control compound, was prepared from nitrobenzodioxolylethanol **8** by dissolution in acetonitrile containing a few drops of water, followed by irradiation with a 400 nm light source for 1 h.

**Scheme 4.**
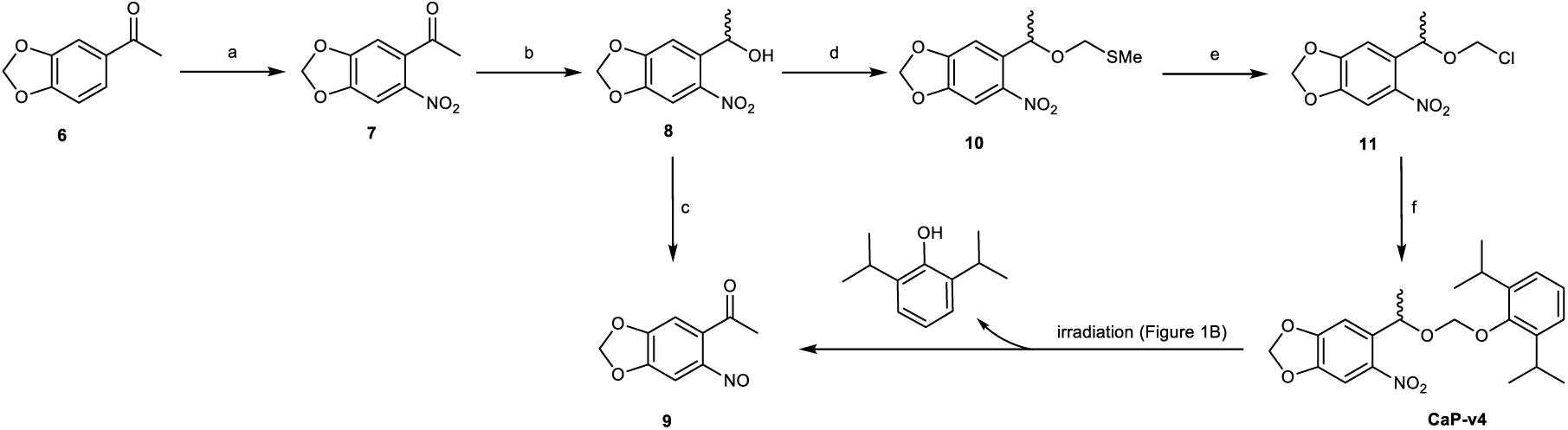
Synthesis of 1-(6-nitrosobenzo[*d*][1,3]dioxol-5-yl)ethan-1-one (**9**) and (*R,S*)-5-(1-((2,6-diisopropylphenoxy)methoxy)ethyl)-6-nitrobenzo[*d*][1,3]dioxole **(CaP-v4).** a. CH_3_NO_2_, HNO_3_, 10 °C, 5 h. b. NaBH_4_, THF, RT, 18 h. c. Irradiation (hϖ =400 nm), 1 h. d. DMSO, Ac_2_O, AcOH, RT, 18 h. e. SO_2_Cl_2_, DCM, 0 °C, 4 h. f. Propofol, NaH, HMPA, RT, 18h.

### Experimental details

#### Chemicals and analytical methods

Chemicals were purchased from Sigma-Aldrich (Darmstadt, Germany), BLD Pharmatech (Kaiserslautern, Germany), Alfa Aesar (Kandel, Germany), abcr (Karlsruhe, Germany), Acros Organics (Geel, Belgium), Thermo Fisher Scientific (Germany) or TCI (Zwijndrecht, Belgium) and used without further purification. Anhydrous solvents were obtained from Acros Organics or dried using molecular sieves following established protocols. Reactions were performed under air unless otherwise stated.

Automated flash chromatography was performed on a Teledyne ISCO CombiFlash Rf 200 (Lincoln, USA). Preparative column chromatography was performed using silica gel (40–63 or 63–200 µm, 60 Å pore size). Ultrasonication was performed in a Bandelin (Berlin, Germany) Sonorex RK 106 35 kHz ultrasonication bath.

The reaction progress was monitored using thin-layer chromatography on Merck Silica gel 60 F254 aluminum plates (Darmstadt, Germany). Detection was performed by UV light at λ = 254 nm or λ = 365 nm, using an Advion Plate Express TLC plate reader coupled with the expression compact mass spectrometer (referred to as TLC-MS) (Ithaca, USA). Melting points (mp) were determined in open capillaries on a Büchi Melting Point apparatus B-545 (Essen, Germany) and are uncorrected.

Mass spectra were recorded on an Agilent (Santa Cruz, USA) InfinityLab LC/MS with Electrospray ionization (ESI) ionization using a Macherey-Nagel EC50/2 Nucleodur C18 Gravity 3 µm column (Düren, Germany) with a flowrate of 0.5 mL/min. The column temperature was 40 °C. Compounds were dissolved in water, methanol, or acetonitrile at a concentration of 1 mg/mL. The injected sample volumes were 3 µL. The following standard gradient was applied: 90% A (water containing 2 mM ammonium acetate) for 10 min, then from 90% A to 100% B (100% acetonitrile) with 10 min followed by flushing with 100% B for 5 min. Positive total ion scans (TIC) were observed from 100-1000 m/z. The UV absorption was detected from λ = 190-900 nm using a diode array detector (DAD). The purity was determined at λ = 220-600 nm.

^1^H NMR (500 MHz) and ^13^C NMR (126 MHz) spectra in DMSO-d_6_ or CDCl_3_ were recorded on a Bruker Avance DRX 500 or Avance Neo 500 (Rheinstetten, Germany), and ^1^H NMR (600 MHz), and ^13^C NMR (151 MHz) spectra were measured on a Bruker Avance III 600 NMR spectrometer at 30 °C. Chemical shifts (δ) are expressed in ppm and referenced to residual solvent peaks of DMSO-d_6_ (2.50/39.52 ppm) or CDCl_3_ (7.26/77.16 ppm). Data are reported as follows: δ, multiplicity (s = singlet, d = doublet, t = triplet, q = quartet, spt = septet and m = multiplet), integration, and coupling constant in Hertz (Hz). Solvent and impurity assignments were based on the data from Fulmer et al.(*47*). Raw NMR data were processed and analyzed using ACD/NMR Processor Academic Edition.

For the uncaging experiments, a 0.5 mg/ml solution of the caged compounds in Hellma quartz cuvettes (type QS-100) (10 mm light path) (Müllheim, Germany) was orthogonally irradiated through the clear sides of the cuvette using a Sahlmann Photochemical Solutions LED controller with LEDs from Nichia, Seoul and Roithner emitting monochromatic wavelengths (Bad Segeberg, Germany): λ = 365 nm (full width at half maximum (FWHM) 9 nm, typical total optical power: 3 x 1050 mW), λ = 400 nm (FWHM 14 nm, typical total optical power: 3 x 1150 mW), λ = 420 nm (FWHM 20 nm, typical total optical power: 3 x 1000 mW), λ = 470 nm (FWHM 30 nm, typical total optical power: 3 x 1250 mW), λ = 525 nm (FWHM 37 nm, typical total optical power: 3 x 500 mW), λ = 590 nm (FWHM 92 nm, typical total optical power: 3 x 525 mW), λ = 690 nm (FWHM 24 nm, typical total optical power: 3 x 450 mW). The progress of uncaging was monitored by changes in the area under the curve (AUC) as determined by HPLC using the standard method described above.

The *cis*/*trans*-isomer ratios of the dark-adapted state (thermodynamic equilibrium) and after successive irradiation for 10 min at different wavelengths (λ = 400 nm and λ = 525 nm) of QAQ and CQAQ (Suppl. Table 2) were determined by integrating the AUC of the respective peaks using HPLC-MS coupled to a diode-array detector (DAD) at the isosbestic point. The photoswitches QAQ and CQAQ were characterized with a ThermoScientific Genesys 50 UV/Vis spectrophotometer (Dreieich, Germany). All measurements were conducted using 50 µM solutions of the respective compound in phosphate-buffered saline (PBS), at pH 7.4. The solutions were stored in the dark for 24 h at room temperature after preparation and prior to measurements. Absorption spectra of the respective photostationary states (PSS) of all compounds were measured in the dark-adapted state (thermodynamic equilibrium) and after irradiation for 1 min at different wavelengths. Photoswitching was determined by repeated alternating irradiation of a single sample for 1 min using the most efficient wavelengths. Thermal relaxation half-lives (t_1/2_) were determined by irradiating the solutions with a previously identified optimal wavelength for 1 min to obtain the thermodynamically unfavored isomer, followed by measuring the absorbance at the respective λ_max_ for a period of 2 h, with intervals of 30 s. The obtained absorption data were fitted to the following equations, which were used to calculate t_1/2_, defined as *t*_1/2_ = *ln*(2)/*k*, using Solver in Microsoft Excel:

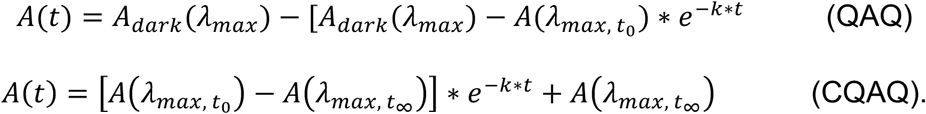

#### Syntheses and analytical data of CaP-v1 to CaP-v4

1-((2,6-Diisopropylphenoxy)methyl)-4,5-dimethoxy-2-nitrobenzene (**CaP-v1**).

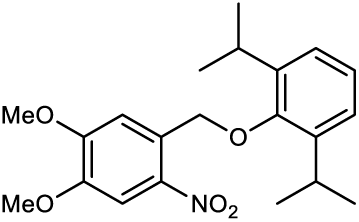

Propofol (50 mg, 0.28 mmol, 1.0 equiv), 1-(bromomethyl)-4,5-dimethoxy-2-nitrobenzene (**1**) (116 mg, 0.42 mmol, 1.5 equiv) and K₂CO₃ (78 mg, 0.56 mmol, 2.0 equiv) were dissolved in dry DMF (1 mL) in an amber glass vial. The reaction mixture was stirred at room temperature for 14 h. The reaction was quenched by the addition of water (2 mL) and the reaction mixture was extracted with ethyl acetate (EtOAc, 20 mL × 3). The combined organic phases were dried over MgSO₄, filtered, and concentrated under reduced pressure. The crude product was purified by column chromatography on silica gel using EtOAc in petroleum ether (gradient 0–20%) to afford the corresponding product. Yield: 46 mg (0.12 mmol, 44%); mp: 80-83 °C; ^1^H NMR (500 MHz, DMSO-d_6_) δ 7.75 (s, 1H), 7.55 (s, 1H), 7.08 - 7.19 (m, 3H), 5.16 (s, 2H), 3.94 (s, 3H), 3.90 (s, 3H), 3.22 (spt, *J* = 6.83 Hz, 2H), 1.17 (d, *J* = 6.94 Hz, 12H); ^13^C NMR (126 MHz, DMSO-d_6_) δ 153.6, 152.7, 147.6, 141.2, 138.6, 128.7, 125.0, 124.0, 109.4, 108.1, 72.4, 56.1, 56.0, 26.2, 23.7; LC-MS (ESI) *m/z* 391.3 ([M+NH_4_]^+^); purity 99% (HPLC-UV, 220-600 nm); HRMS (ESI) (*m/z*): [M+Na]^+^ calculated for C_12_H_27_NNaO_5_, 396.1781; found, 396.1796.

2,6-Diisopropylphenyl (4,5-dimethoxy-2-nitrobenzyl)carbonate (**CaP-v2**).

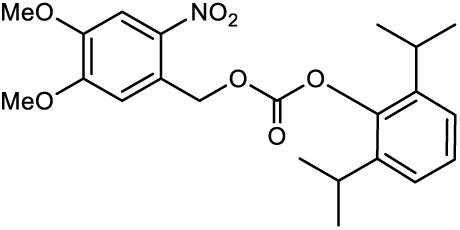

4,5-Dimethoxy-2-nitrobenzyl carbonochloridate (**3**)(*44*).

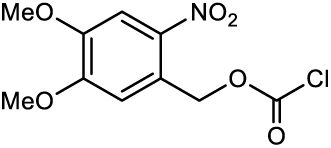

(4,5-Dimethoxy-2-nitrophenyl)methanol (**2**) (500 mg, 2.3 mmol, 1 equiv), and triphosgene (740 mg, 2.3 mmol, 1 equiv), were dissolved in dry tetrahydrofuran (THF, 7 ml) and the mixture was cooled to 0°C in an ice bath and stirred. Then, a solution of Et_3_N (300 µl, 2.3 mmol, 1 equiv) in dry THF (7 ml) was added dropwise. The reaction was allowed to warm to room temperature and stirred for 18 h. The mixture was then filtered, and concentrated in vacuo. Product **3** was recrystallized from toluene. Yield: 253 mg, (0.92 mmol, 40%); ^1^H NMR (500 MHz, DMSO-d_6_) δ 7.68 (s, 1H), 7.35 (s, 1H), 5.03 (s, 2H), 3.91 (s, 3H), 3.88 (s, 3H); ^13^C NMR (126 MHz, DMSO-d_6_) δ 153.14, 148.88, 140.50, 126.74, 114.52, 108.89, 56.69, 56.47, 43.56.

A stirred solution of propofol (50 mg, 0.28 mmol, 1.0 equiv) in dry dichloromethane (DCM, 5 ml) was cooled to 0°C in an ice bath. Carbonochloridate (**3**, 150 mg, 0.56 mmol, 2.0 equiv) and Et_3_N (120 µl, 0.84 mmol, 3.0 equiv) were then added, and the solution was stirred for 2.5 h. When no further reaction progress was detected, the mixture was quenched with water (10 mL) and extracted with DCM (3 x 20 mL). The combined organic phases were dried over MgSO_4_, filtered and concentrated in vacuo. The crude product was purified by column chromatography on silica gel using EtOAc in petroleum ether (30%) to afford the desired product **Cap-v2**. Yield: 48 mg (0.12 mmol, 41%). mp: 106-108 °C. ^1^H NMR (500 MHz, DMSO-d_6_) δ 7.73 (s, 1H), 7.27 (s, 1H), 7.18 - 7.26 (m, 3H), 5.58 (s, 2H), 3.90 (s, 3H), 3.89 (s, 3H), 2.94 (quin, *J* = 6.86 Hz, 2H), 1.13 (d, *J* = 6.94 Hz, 12H);^13^C NMR (126 MHz, DMSO-d_6_) δ 153.0, 152.8, 148.5, 145.2, 140.3, 139.9, 126.9, 124.5, 124.1, 112.4, 108.4, 66.9, 56.2, 56.1, 26.7, 22.9. LC-MS (ESI) *m/z* 435.4 ([M+NH_4_]^+^); purity 98% (HPLC-UV (220-600 nm); HRMS (ESI) (*m/z*): [M+NH_4_]^+^ calculated for C_22_H_31_N_2_O_5_, 435.2126; found, 435.2102.

1-(((2,6-Diisopropylphenoxy)methoxy)methyl)-4,5-dimethoxy-2-nitrobenzene (**CaP-v3**).

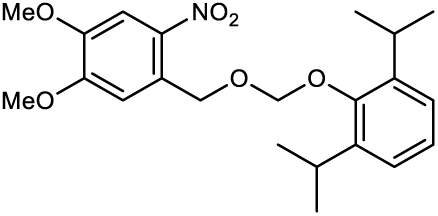

((2,6-Diisopropylphenoxy)methyl)(methyl)sulfane (**4**) (*45*):

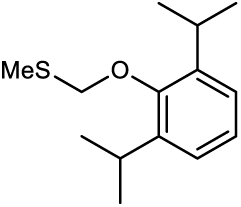

Under an argon atmosphere, sodium hydride (60% dispersion in mineral oil, 250 mg, 6.30 mmol, 1.6 equiv) was suspended in 10 mL of HMPA and stirred. To this suspension, propofol (660 µL, 720 mg, 4.04 mmol, 1 equiv) was added dropwise over 15 min. The reaction was stirred at room temperature for 30 min. Subsequently, chloromethyl methyl sulfide (541 µL, 624 mg, 6.46 mmol, 1.6 equiv) was added dropwise, and stirring continued at room temperature for 18 h. The reaction was quenched by the addition of water (30 mL) and stirred for 10 min. The mixture was extracted with EtOAc (3 x 40 mL), and the combined organic phases were washed twice with water (10 mL each), dried over MgSO_4_, filtered, and concentrated under reduced pressure. The crude oily residue was purified by flash chromatography on silica gel, using petroleum ether and EtOAc, applying a gradient (0 to 5%), yielding 460 mg (1.93 mmol, 50%) of the desired compound **4** as a colorless oil. ^1^H NMR (500 MHz, CDCl_3_) δ 7.13 (s, 3H), 4.88 (s, 2H), 3.39 (spt, *J* = 6.88 Hz, 2H), 2.38 (s, 3H), 1.25 (d, *J* = 6.97 Hz, 12H). LC-MS (ESI) *m/z* 237.1 ([M - H]^-^); purity 97% (HPLC-UV, 220-600 nm). 2-(Chloromethoxy)-1,3-diisopropylbenzene (**5**) (*45*):

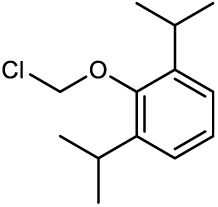

A solution of methylsulfane (**4**, 660 mg, 2.77 mmol, 1 equiv) in dry DCM (5 mL) was stirred under an argon atmosphere. While maintaining a temperature of 0 °C in an ice bath, a 1 M solution of SO₂Cl₂ in dry DCM (3.1 mL, 3.1 mmol, 1.1 equiv) was added dropwise. The reaction was kept at 0 °C for 30 min, followed by stirring at ambient temperature for an additional 3 h. After completion, the solvent was removed under reduced pressure affording 439 mg (1.94 mmol, 70%) of the desired compound **5** as a colorless oil which was used for the next step without further purification. ^1^H NMR (600 MHz, CDCl_3_) δ 7.12 - 7.22 (m, 3H), 5.77 (s, 2H), 3.36 (spt, *J* = 6.88 Hz, 2H), 1.24 (d, *J* = 6.79 Hz, 12H). LC-MS (ESI) *m/z* 249 ([M + Na]^+^); purity 90% (HPLC-UV, 220-600 nm).

Sodium hydride (60% dispersion in mineral oil, 20 mg, 0.47 mmol, 1 equiv) was suspended in HMPA (2 mL) upon stirring under an inert argon atmosphere. 2-Nitrophenylmethanol (**2**,100 mg, 0.47 mmol, 1 equiv) was added portionwise, and the mixture was stirred at RT for 1 h. Subsequently, diisopropylbenzene (**5**, 117 mg, 0.52 mmol, 1.1 equiv) was dissolved in 1 mL HMPA and added dropwise to the reaction mixture; then stirring was continued at room temperature for 18 h. The reaction mixture was diluted with water (3 mL) upon stirring. The aqueous phase was separated and extracted twice with EtOAc (50 mL each). The combined organic phases were washed twice with water (50 mL each), dried over anhydrous magnesium sulfate, filtered, and concentrated under reduced pressure. The crude residue was purified by flash column chromatography using petroleum ether and EtOAc applying a gradient (0 to 2%) followed by purification using preparative TLC with cyclohexane and EtOAc (99:1) to afford 3 mg (0.007 mmol, 2%) of the desired product **CaP-v3** as a colorless oily substance. ^1^H NMR (500 MHz, DMSO-d_6_) δ 7.71 (s, 1H), 7.27 (s, 1H), 7.07 - 7.15 (m, 3H), 5.17 (s, 2H), 5.12 (s, 2H), 3.83 - 3.91 (m, 6H), 3.25 - 3.30 (m, 2H), 1.13 (d, J = 6.84 Hz, 12H); ^13^C NMR (126 MHz, DMSO-d_6_) δ 153.3, 151.5, 147.5, 141.2, 139.2, 128.7, 124.9, 123.9, 110.2, 108.1, 98.5, 67.7, 56.1, 56.0, 26.2, 23.7. LC-MS (ESI) *m/z* 421.3([M+NH_4_]^+^); purity 95% (HPLC-UV (220-600 nm), HRMS (ESI) (*m/z*): [M+Na]^+^ calculated for C_22_H_29_NNaO_6_, 426.1887; found, 426.1898.

(*R,S*)-5-(1-((2,6-diisopropylphenoxy)methoxy)ethyl)-6-nitrobenzo[*d*][1,3]dioxole (**CaP-v4, ‘CaP’**).

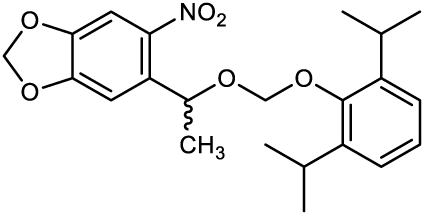

1-(6-Nitrobenzo[*d*][1,3]dioxol-5-yl)ethan-1-one (**7**) (*46*):

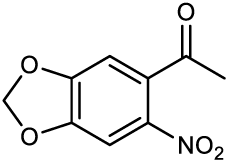

A solution of benzodioxolylethenone (**6**, 10 g, 0.6 mol, 1 equiv) in nitromethane (CH_3_NO_2_) (111 mL, 126.42 g, 2.07 mol, 3.45 equiv) was prepared and stirred in an ice-water mixture. Concentrated nitric acid (HNO_3_) (22 mL, 0.49 mol, 0.8 equiv) was then added dropwise using a dropping funnel while maintaining continuous stirring and avoiding an increase in temperature above 10 °C by adding ice to the water bath. Then, the mixture was kept at room temperature and stirred for an additional 5 h, while controlling the reaction progress by TLC. After consumption of the starting material, the reaction was quenched by slowly adding a saturated aqueous sodium hydrogencarbonate (NaHCO_3_) solution (500 mL, pH 9) upon stirring using a dropping funnel. The resulting mixture was transferred to a separatory funnel and extracted three times with EtOAc (250 mL each). The combined organic phases were washed with brine, dried over MgSO_4_, filtered, and concentrated under reduced pressure. The crude residue was purified by column chromatography on silica gel, eluting with petroleum ether and EtOAc (4:1), affording compound **7** as a light yellow solid (5.9 g, 0.03 mol, 46% yield). ^1^H NMR (500 MHz, DMSO-d_6_) δ 7.68 (s, 1H), 7.29 (s, 1H), 6.29 (s, 2H), 2.49 (s, 3H); ^13^C NMR (126 MHz, DMSO-d_6_) δ 198.6, 152.0, 148.8, 140.2, 133.2, 106.7, 104.6, 104.0, 29.7. LC-MS (ESI) *m/z* 210.2 ([M + H]^+^); purity 90% (HPLC-UV, 220-600 nm).

(*R,S*)-1-(6-Nitrobenzo[*d*][1,3]dioxol-5-yl)ethan-1-ol (**8**) (*46*):

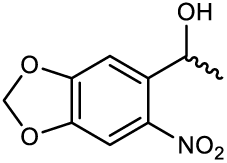

Nitrobenzodioxolylethenone (**7**, 4.9 g, 23.43 mmol, 1 equiv) was dissolved in anhydrous THF (115 mL), and NaBH_4_ (1.77 g, 46.85 mmol, 2 equiv) was added. The reaction was stirred at room temperature for 18 h and then quenched with 2N aqueous HCl solution until the formation of gas stopped. The mixture was poured into brine (100 mL) and the organic components were extracted with EtOAc three times (100 mL each). The combined organic phases were washed with brine (100 mL), dried over anhydrous MgSO_4_, filtered, and concentrated under reduced pressure. The crude residue was purified by flash column chromatography using a mixture of cyclohexane and EtOAc, applying a gradient (1 to 40%), affording compound **8** as a yellow solid (3.41 g, 16.15 mmol, 69% yield). ^1^H NMR (600 MHz, CDCl_3_) δ 7.46 (s, 1H), 7.28 (s, 1H), 6.12 (d, *J* = 1.10 Hz, 1H), 6.12 (d, *J* = 1.10 Hz, 1H), 5.46 (q, *J* = 6.36 Hz, 1H), 1.54 (d, *J* = 6.24 Hz, 3H). LC-MS (ESI) *m/z* 229.2 ([M + NH_4_]^+^); purity 90% (HPLC-UV, 220-600 nm).

1-(6-Nitrosobenzo[*d*][1,3]dioxol-5-yl)ethan-1-one (**9**):

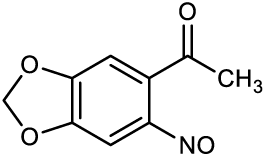

Nitrobenzodioxolylethanol (**8**, 100 mg, 0.47 mmol, 1 equiv) was dissolved in acetonitrile (10 mL), containing few drops of water, in a round-bottom flask. The mixture was irradiated with a 400 nm light source for 1 h. After completion of the photoreaction, the solvent was removed under reduced pressure. The resulting crude product was purified by flash column chromatography on silica gel, eluting with a gradient of EtOAc in cyclohexane (3% to 50%), yielding the desired compound **9** as a green solid (26 mg, 0.13 mmol, 28% yield). mp: 112-114 °C. ^1^H NMR (500 MHz, DMSO-d_6_) δ 7.42 (s, 1H), 6.27 (s, 2H), 6.23 (s, 1H), 2.75 (s, 3H); ^13^C NMR (126 MHz, DMSO-d_6_) δ 201.5, 160.8, 154.4, 150.1, 143.3, 106.6, 104.0, 88.8, 32.9. LC-MS (ESI) *m/z* 194.0 ([M + H]^+^); purity 99% (HPLC-UV, 220-600 nm).

(*R,S*)-5-(1-((methylthio)methoxy)ethyl)-6-nitrobenzo[*d*][1,3]dioxole (**10**) (*46*):

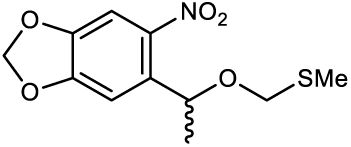

To a solution of nitrobenzodioxolylethanol (**8**, 458 mg, 2.17 mmol, 1 equiv) in DMSO (20 mL), acetic anhydride (15 mL) and acetic acid (10 mL) was added and the mixture was stirred at room temperature for 18 h. Upon completion, the reaction mixture was added dropwise to a saturated aqueous solution of sodium hydrogen carbonate (50 mL) upon stirring, and it was allowed to stand for 18 h. The resulting mixture was then extracted four times with EtOAc (50 mL each). The combined organic phases were dried over anhydrous MgSO_4_, filtered, and concentrated under reduced pressure. The crude product was purified by flash column chromatography on silica gel, eluting with a gradient of EtOAc in petroleum ether (0% to 50%), yielding the product **10** as a light yellow semisolid product (230 mg, 0.85 mmol, 39% yield). ^1^H NMR (500 MHz, DMSO-d_6_) δ 7.56 (s, 1H), 7.15 (s, 1H), 6.22 (dd, *J* = 0.86, 8.35 Hz, 2H), 5.26 (q, *J* = 6.31 Hz, 1H), 4.64 (d, *J* = 11.44 Hz, 1H), 4.31 (d, *J* = 11.58 Hz, 1H), 2.03 (s, 3H), 1.42 (d, *J* = 6.31 Hz, 3H); ^13^C NMR (126 MHz, DMSO-d_6_) δ 152.1, 146.9, 141.9, 135.7, 106.0, 104.5, 103.3, 72.4, 69.4, 22.9, 13.3. LC-MS (ESI) *m/z* 272.3 ([M + H]^+^); purity 99% (HPLC-UV, 220-600 nm).

(*R,S*)-5-(1-(chloromethoxy)ethyl)-6-nitrobenzo[*d*][1,3]dioxole (**11**) (*46*):

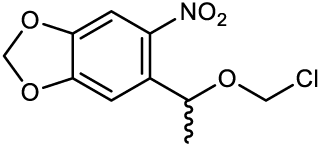

A 1 M solution of sulfuryl chloride (5.1 mL, 5.1 mmol, 6.6 equiv) in DCM was added dropwise to a stirred solution of nitrobenzodioxole (**10**, 210 mg, 0.77 mmol, 1 equiv) in dry DCM (5 mL) in an ice bath. The reaction mixture was then stirred for 4 h at 0 °C. The solvent and volatile by-products were subsequently removed under reduced pressure. Chloroform (5 mL) was added to the residue and evaporated under reduced pressure; this process was repeated once more. The resulting yellowish residue of compound **11** was used directly in the subsequent step without further purification (192.25 mg, 0.74 mmol, 96% yield). ^1^H NMR (600 MHz, CDCl_3_) δ 7.48 (s, 1H), 7.06 (s, 1H), 6.10 (d, *J* = 4.59 Hz, 3H), 5.54 (q, *J* = 6.36 Hz, 1H), 5.49 (d, *J* = 6.05 Hz, 1H), 5.22 (d, *J* = 6.05 Hz, 1H), 1.54 (d, *J* = 6.42 Hz, 4H).

(*R,S*)-5-(1-((2,6-diisopropylphenoxy)methoxy)ethyl)-6-nitrobenzo[*d*][1,3]dioxole (**CaP-v4, ‘CaP’**).

NaH (30 mg, 0.74 mmol, 60% dispersion in mineral oil, 1.1 equiv) was suspended in HMPA (2.5 mL) under an argon atmosphere and stirred. Propofol (120 mg, 0.67 mmol, 1 equiv) was then added to the suspension, and the reaction mixture was stirred at room temperature for an additional 30 min. Nitrobenzodioxole (**11**, 192.25 mg, 0.74 mmol, 1.1 equiv) was then added, and stirring was continued for 18 h at room temperature. The reaction was quenched by the addition of water (5 mL), and stirring was maintained for 10 min. The aqueous mixture was extracted with EtOAc (3 × 50 mL), and the combined organic phases were washed twice with water (20 mL each), dried over anhydrous MgSO_4_, filtered, and concentrated under reduced pressure. The crude residue was purified by flash column chromatography on silica gel, using a gradient of EtOAc (0–20%) in DCM, affording the target compound **CaP-v4 (‘CaP’)**.

Recrystallization was performed by dissolution in methanol, followed by sonication using an ultrasound bath to precipitate a white solid that was filtered off and dried under high vacuum (42 mg, 0.10 mmol, 16%). mp: 109-111 °C. ^1^H NMR (600 MHz, DMSO-d_6_) δ 7.61 (s, 1H), 7.16 (s, 1H), 7.07 (s, 3H), 6.18 - 6.26 (m, 2H), 5.45 (q, *J* = 6.24 Hz, 1H), 4.89 (d, *J* = 5.87 Hz, 1H), 4.75 (d, *J* = 5.87 Hz, 1H), 3.13 (spt, *J* = 6.88 Hz, 2H), 1.51 (d, *J* = 6.24 Hz, 3H), 1.09 (d, *J* = 6.79 Hz, 6H), 1.05 (d, *J* = 6.79 Hz, 6H); ^13^C NMR (151 MHz, DMSO-d_6_) δ 152.2, 150.7, 147.0, 141.6, 141.2, 135.8, 124.9, 123.8, 106.2, 104.5, 103.4, 96.1, 71.1, 26.1, 23.7, 23.5, 22.9. LC-MS (ESI) *m/z* 419.2 ([M + NH_4_]^+^); purity 99% (HPLC-UV, 220-600 nm); HRMS (ESI) (*m/z*): [M+Na]^+^ calculated for C_22_H_27_NNaO_6_, 424.1731; found, 424.1731.

### Electrophysiological recordings and stimulation

#### Slice preparation for patch clamp recordings

To examine the effect of the photoactivatable compounds on neuronal activity in vitro, adult wild type mice were sacrificed by decapitation and brains were extracted. Postsurgical human brain tissue was collected in ice-cold carbogenated (95 % O2 and 5 % CO2) sucrose artificial cerebrospinal fluid (sACSF (mM): 60 NaCl, 100 Sucrose, 2.5 KCl, 1.25 NaH_2_PO_4_, 26 NaHCO_3_, 1 CaCl_2_, 5 MgCl_2_, 20 Glucose; pH 7.4; 300-310 mOsm) from the operation room right after resection. For both murine and human tissue, coronal brain sections were sliced with a vibratome (Leica VT1200S vibratome platform) in ice-cold carbogenated sACSF at a thickness of 300µm (mouse tissue) or 400µm (human tissue). After cutting, murine and human slices were incubated at 35 °C in carbogenated sACSF for 30 min before being transferred to carbogenated artificial cerebrospinal fluid (ACSF; mM: 125 NaCl, 3 KCl, 1.25 NaH_2_PO_4_, 26 NaHCO_3_, 2.6 CaCl_2_, 1.3 MgCl_2_, 15 Glucose; pH 7.4; 300-310 mOsm) at room temperature (RT).

#### Patch clamp recordings

Whole-cell patch clamp experiments were conducted at 35 °C (heat controller: Luigs and Neumann temperature controller V) in slices perfused with ACSF (125 – 200 ml/hour) in a submerged recording chamber. Slices were fixed in place with a ‘harp’ made from dental floss glued over a platinum frame. Recording pipettes (resistance 5-8 MΩ) were made from borosilicate glass capillaries (Science products, GB150F-8P) using an upright puller (Narishige Japan, Model PP-830). For visualization of CA1 pyramidal neurons, a fixed stage upright microscope (Olympus BX51WI) with a 60x water immersion objective (Olympus LUMPlanFl 60x/0.90 W ∞/0 IR) was used. Signals were amplified (x10, Cornerstone by Dagan, BVC-700A; 10 kHz low-pass filter), digitized at 10 – 100 kHz (Axon Digidata 1440A) and recorded using the software Clampex 10.7 (Molecular Devices). Neurons showing resting membrane potentials above −50 mV (and −40 mV at the end of the recording) were excluded from the experiment.

##### QAQ and CQAQ

Action potential (AP) recordings in CA1 neurons were performed in current clamp configuration at a holding potential of −70 mV. K+-based intracellular solution used for filling the patch pipettes contained (mM): 115 K-gluconate, 20 KCl, 10 Na-phosphocreatine, 10 HEPES, 2 Mg-ATP, 0.3 Na-GTP; or 110 K-gluconate, 1 CaCl_2_*2H_2_O, 10 Na-phosphocreatine, 10 HEPES, 11 EGTA, 0.3 Na-GTP; pH 7.2; ∼290 mOsm). The internal liquid junction potential was −14.6 or −15.9 mV. Photoswitchable lidocaine derivatives QAQ (QAQ dichloride, Tocris Bioscience) and CQAQ (Trads et al., 2019) were first dissolved in distilled water (QAQ: 25 mM; CQAQ: 5 mM), further dissolved in the intracellular solution to a final concentration of 100 µM, and applied internally to the cell through the patch pipette. To allow equilibration time, measurements were started 5-10 min after patching a neuron. APs were evoked with a depolarizing square pulse applied through the patch pipette. Each stimulation protocol consisted of three sweeps of square pulse stimulation (pulse duration: 1s; current steps: −50 to −250 pA, 2x rheobase, 3x rheobase; rheobase: 30 – 250 pA). After stable AP activity was recorded, the PD was first photo-inactivated, subsequently photoactivated and finally inactivated again. The stimulation protocol was repeated for every photoactivation or –inactivation step. For photostimulation of QAQ, illumination at 525±35 nm (“trans-activation”; power at the sample: 1.08 mW / mm^2^; Thorlabs) or 387±5.5 nm (“cis-inactivation”; power at the sample: 0.35 mW / mm^2^; Thorlabs) was applied in the entire field of view (FOV) by a halogen lamp (X-Cite 120 –Fluorescence Illumination System) for 1 min each (30 s prior to recording plus 30 s recording time). The sign-inverted CQAQ compound was activated at 400±20 nm (power at the sample: 1.53 mW / mm^2^; filter: FBH400-40, Thorlabs) and inactivated at 525 nm. Light power was measured with a photometer (Optical Power Meter, PM400, Thorlabs GmbH) and the sensor was placed at the same distance to the objective as the tissue sample. Negative control experiments were conducted in an identical fashion to the stimulation protocol described above with the only difference that PDs were not added to the intracellular solution. Neurons showing a change of input resistance of > 30 % during the experiment (input resistance: 201.51 ± 9.57 MΩ, SEM [mouse tissue], 260.36 ± 24.63 MΩ, SEM [human tissue]) were excluded from the analysis.

##### Caged Propofol (CaP)

Inhibitory postsynaptic currents (IPSCs) of CA1 neurons were recorded in voltage clamp configuration at 0 mV. Cs^+^-based intracellular solution used for filling patch pipettes contained (mM): 120 Cs methanesulfonate, 0.5 MgCl2, 5 HEPES, 5 EGTA, 5 Na2-ATP, 5 QX-314 Cl^-^; pH 7.4; 282 mOsm. Cl^-^ reversal potential: −80.56 mV. The internal liquid junction potential was −20 mV. CaP was first dissolved in dimethyl sulfoxide (DMSO, Sigma-Aldrich Chemie GmbH; 100 mM), further dissolved in ACSF to a final concentration of 100 µM (final DMSO concentration: 0.5 %), and applied to the recording chamber by external perfusion. IPSCs were elicited by stimulation of the Schaffer Collaterals using a bipolar electrode (30209 Cluster 20 Bipolar Electrodes, FHC Inc, USA) controlled by an isolated pulse stimulator (Model 2100 Isolated Pulse Stimulator, A-M systems). Each stimulation protocol consisted of three sweeps of square pulse stimulation (pulse duration: 0.2 ms; current: 100 – 530 µA). After stable baseline IPSC activity was recorded, CaP/ACSF solution was washed into the recording chamber and the stimulation protocol was repeated. For uncaging of CaP, illumination at 400±20 nm (uncaging time: 3 min; power: 1.53 mW / mm^2^; filter: FBH400-40, Thorlabs) was applied in the entire FOV using a halogen lamp. Subsequently uncaged Propofol was washed out of the recording chamber with CaP/ACSF solution. Positive control experiments were performed with commercially available propofol (100 μM, pre-dissolved in DMSO [final DMSO concentration: 0.5%]; propofol 1 % [10 mg / 1 ml] MCT Fresenius G-Amp, Fresenius Kabi Deutschland GmbH). Negative control experiments were conducted with the end product of the photoprotective group (ePPG) in ACSF (100 µM, pre-dissolved in DMSO [final DMSO concentration: 0.5 %]).

### Slice preparation for recording epileptiform activity (EA)

To investigate the effect of the photoactivatable compounds on EA in vitro, adult wild type mice were sacrificed by decapitation and their brains were extracted. Postsurgical human brain tissue was collected in ice-cold carbogenated sACSF from the operation room right after resection. Pia and blood vessels were removed from human cortical tissue blocks and the blocks were cut vertical to the cortical surface. Mouse (horizontal) or human cortical slices were cut with a vibratome in ice-cold carbogenated sACSF at a thickness of 400 µm and transferred to carbogenated ACSF (RT).

#### Induction of EA

Prior to recording local field potentials (LFP) in neocortex, slices were perfused (150 ml/hour) with ACSF in an interface slice chamber (BSC2-2, World Precision Instruments) at 32 °C (mouse tissue) or 35 °C (human tissue) (temperature controller: PTC-03, World Precision Instruments) for at least one hour to allow sufficient tissue oxygenation. LFP recordings were performed in cortical slices in an interface chamber (BSC1, World Precision Instruments) perfused (250 ml/hour) with modified ACSF (mACSF) containing no Mg^2+^ (mM: 129.1 NaCl, 3 KCl, 1.25 NaH_2_PO_4_, 26 NaHCO_3_, 1.6 CaCl_2_, 15 Glucose; pH 7.4; 300-310 mOsm) at 36°C (temperature controller: PTC03, World Precision Instruments). Slice incubation in the interface chamber and with Mg^2+-^free ACSF reliably induced interictal epileptic activity (IEA; incubation time ranging from approximately 2 – 20 min [mouse tissue] or 30 – 90 min [human tissue]) and seizure like episodes (SLE; occurred in some slices; in average SLE started 3.35 min [mouse tissue] or 6 min [human tissue] after IEA) in mouse and human cortical slices.

#### Local field potential (LFP) recordings

Recording pipettes (resistance 5 - 8 MΩ) were pulled using a vertical puller (Narishige Japan, Model PP-830) from borosilicate glass capillaries (Science products, GB150F-8P). For slice visualization, a commercially available stereomicroscope (Leica A60 F, Leica Microsystems GmbH) was used. Signals were amplified (x100; Cornerstone by Dagan, BVC-700A; 1 kHz low-pass-filter), digitized at 10 – 100 kHz (Axon Digidata 1440A) and recorded using the Clampex 10.7 software (Molecular Devices). 50 Hz noise was eliminated by a HumBug (Digitimer). ACSF was used for filling patch pipettes. CaP was first dissolved in DMSO (90 mM), further dissolved in mACSF to a final concentration of 90 µM (final DMSO concentration: 0.5 %), and applied to the recording chamber by external perfusion. LFP recording were conducted in gap-free recording mode. After stable baseline EA was recorded, CaP/mACSF solution was washed into the recording chamber. To allow sufficient distribution time, CaP was uncaged after 4 min of wash in. For uncaging of CaP, illumination at 400 nm (uncaging time: 6 min; power: 0.5 mW / mm^2^) was applied in the entire FOV (Spectra Light Engine, Lumencor Inc.). Subsequently uncaged Propofol was washed out with CaP/mACSF solution. Positive control experiments were performed using commercially available propofol (100 μM, pre-dissolved in DMSO [final DMSO concentration: 0.5 %]; propofol 1 % [10 mg / 1 ml] MCT Fresenius G-Amp, Fresenius Kabi Deutschland GmbH). Negative control experiments were conducted with the ePPG in mACSF (100 µM, pre-dissolved in DMSO [final DMSO concentration: 0.5 %]).

### Immunohistochemistry

#### NeuN staining

Immunhistochemical NeuN staining was performed in a similar fashion as previously described (Mitlasóczki et al., 2025). Mice were deeply anesthetized with 16 mg / kg xylazine (20 mg / mL, i.p., xylazine hydrochloride, Serumwerk Bernburg AG) and 100 mg / kg ketamine (10 %, Serumwerk Bernburg AG), and transcardially perfused with phosphate-buffered saline (PBS) and 4 % formaldehyde (FA, in PBS). Mouse brains or postsurgical human tissue slices were stored overnight in 4 % FA at 4 °C. Hippocampal or cortical sections (mouse tissue: 40 μm; human tissue: 400 µm) were used for immunofluorescent stainings against NeuN (Synaptic Systems GmbH, 266006, 1:500). Briefly, slices were washed twice in PBS and once in PBS containing 0.25 % Triton (PBT) for 10 min (human tissue: overnight at 4 °C), subsequently blocked for 1 h at RT with PBT containing 10 % normal goat (NGS) serum, and then incubated with the primary antibody diluted in 3 % serum/PBT overnight at 4 °C. After washing three times with PBT for 10 min, slices were incubated with the respective secondary antibody (AlexaFluor488-conjugated goat anti-Chicken, Invitrogen, A32931) for 1.5 h at RT in 3 % serum/PBT, washed twice with PBT for 10 min at RT, and mounted with Aqua-Poly/Mount (Polysciences). Human slices were incubated in PBT only once (overnight), all other washing steps and incubations were performed with PBS. For image acquisition, a Visiscope confocal microscope (Visitron Systems GmbH) attached to a spinning disc unit CSU-W1 (Yokogawa) was used. Images were acquired using a pco.edge sCMOS camera (Excelitas Technologies) and visualized with the VisiView®-Software (Visitron Systems GmbH). Acquired images were further analyzed using ImageJ (NIH).

#### Biocytin Staining

Some patched neurons were stained for Biocytin enabling visualization of neuron morphology and location. By adding Biocytin (3 mg / 1 mL, TRC Canada, B388100) into the intracellular solution, Biocytin diffused into the patched neurons. After the patch clamp recording, slices were fixed in 4 % FA in PBS overnight at 4°C. Slices were washed with PBS for 10 min, and subsequently permeabilized in PBT overnight at 4°C. After washing for 10 min with PBS, slices were incubated in streptavidin coupled to Alexa555 (1:500, Invitrogen) in PBS for 5 min. Slices were washed with PBS for 10 min and incubated in 4′,6-Diamidin-2-phenylindol (DAPI, 1:10000, Biotium) in PBS for 5 min. After washing two times for 10 min with PBS slices were mounted with Aqua-Poly/Mount (Polysciences). Unless stated otherwise, staining steps were performed on a shaker at RT.

### Data analysis

Data from electrophysiological experiments were analyzed using custom code written in Matlab R2023b (MathWorks) or PyCharm 2023.1.2 (JetBrains).

#### AP frequency

Recordings with AP amplitude <40 mV were excluded. Recordings with a shaky or irregular baseline were excluded. APs were detected using the peak-find function with a manually set threshold in Matlab.

#### IPSC decay time and leak current

Recordings with an access resistance variation of >20 % were excluded and data that was averaged over three traces. Decay time was defined as the time between IPSC maximum and the time until the IPSC reaced 37 % of its amplitude(*48*). Leak current was calculated from the difference in membrane current during a −10 mV hyperpolarizing potential step (100 ms; square pulse) prior to IPSC induction.

#### EA characterization and analysis

For detection of EA, signals were high-pass filtered at 0.5 Hz. Threshold for detecting EA was set at ≥3:1 signal to noise ratio and combined with a peak-find function. To be categorized as epileptiform, events had to contain spikes with a half width of <400 ms. Detected EA was double-checked by visual inspection to prevent detection of artifacts. SLE were defined as episodic and rhythmic activity with a minimum duration of 5 s and minimum frequency of 1 Hz (mouse tissue) or 0.5 Hz (human tissue). IEA was defined as spikes or bursts with a rate of ≥5 events / min (mouse tissue) and ≥1 event / min (human tissue) prior to drug wash-in. This minimum occurrence threshold was set to ensure robust quantification of EA.

### Statistical analysis

Statistical analysis was performed with Prism 10.3 (GraphPad), and differences were considered significant at the following p-values: not significant (n.s.) ≥0.05, *<0.05, **<0.01 and ***<0.001, ****<0.0001. For small datasets (e.g. due to limited human tissue availability), a normal distribution was assumed. When comparing two groups, drug effects were tested using a paired t-test. Drug effects in datasets with more than two groups were tested using repeated measures ANOVA followed by Tukey post-hoc test, unless stated otherwise.

## Acknowledgements.

We sincerely thank Lea Adenauer, Laura Kück and Nele Neumann for outstanding technical support. We are grateful to members of the Wenzel, Müller, Trauner, Huberfeld and Beck laboratories for comments. This work was supported by the TRA Life and Health at University of Bonn (M.W., C.E.M.: A-397.0005), and the European Research Council (M.W.: StG #101039945). C.E.M. and F.S. were supported by the Deutsche Forschungsgemeinschaft (DFG, Research Training Group GRK 2873, #494832089), and by the Bonn International Graduate School of Drug Sciences (BIGS-DrugS). D.T. thanks the National Institutes of Health for financial support (Grant R01 GM126228). T.Ke. was supported by DFG Research group FOR-2715. H.B. was supported by DFG grant BE 1822/11-2. M.S. received funding from the Mildred Scheel School of Oncology (German Cancer Aid; #70113307) and the Ministry of Culture and Science of the State of North Rhine-Westphalia as part of the CANTAR research network (#NW21-062H). We also acknowledge the Imaging Core Facility of the Bonn Technology Campus Life Sciences funded by the DFG (project #388169927).

## Author contributions

M.W. conceived the project. M.W., M.B., C.E.M. and A.E. wrote the paper with input from all authors. Experimental contributions: Molecule design, synthesis, purification, and chemical analyses: A.E., C.E.M., T.Ko., D.T. and F.S.; Neurophysiology experiments (patch-clamp, LFP, photo-activation, in vitro models of epileptiform activity, IHC): M.B., M.M.H.N., M.S.S., T.O. and T.Ke.; Data processing and analyses were carried out by M.B., M.W., A.E., M.M.H.N., M.S.S. and F.S.; M.W., C.E.M., H.B., G.H. and R.S. provided infrastructure, experimental, analytical, and clinical expertise.

## References

1. P. Agostinis, K. Berg, K. A. Cengel, T. H. Foster, A. W. Girotti, S. O. Gollnick, S. M. Hahn, M. R. Hamblin, A. Juzeniene, D. Kessel, M. Korbelik, J. Moan, P. Mroz, D. Nowis, J. Piette, B. C. Wilson, J. Golab, Photodynamic therapy of cancer: An update. CA. Cancer J. Clin. 61, 250–281 (2011).

2. L. A. Stokowski, Fundamentals of phototherapy for neonatal jaundice. Adv. Neonatal care. 11, 10–21 (2011).

3. K. Hüll, J. Morstein, D. Trauner, In Vivo Photopharmacology. Chem. Rev. 118, 10710–10747 (2018).

4. B. Blanco, K. A. Palasis, A. Adwal, D. F. Callen, A. D. Abell, Azobenzene-containing photoswitchable proteasome inhibitors with selective activity and cellular toxicity. Bioorganic Med. Chem. 25, 5050–5054 (2017).

5. D. L. Fortin, M. R. Banghart, T. W. Dunn, K. Borges, D. A. Wagenaar, Q. Gaudry, M. H. Karakossian, T. S. Otis, W. B. Kristan, D. Trauner, R. H. Kramer, Photochemical control of endogenous ion channels and cellular excitability. Nat. Methods. 5, 331–338 (2008).

6. A. Mourot, T. Fehrentz, Y. Le Feuvre, C. M. Smith, C. Herold, D. Dalkara, F. Nagy, D. Trauner, R. H. Kramer, Rapid optical control of nociception with an ion-channel photoswitch. Nat. Methods. 9, 396–402 (2012).

7. L. Camerin, G. Maleeva, A. M. J. Gomila, I. Suárez-Pereira, C. Matera, D. Prischich, E. Opar, F. Riefolo, E. Berrocoso, P. Gorostiza, Photoswitchable Carbamazepine Analogs for Non-Invasive Neuroinhibition In Vivo. Angew. Chemie - Int. Ed. 63, e202403636 (2024).

8. A. Polosukhina, J. Litt, I. Tochitsky, J. Nemargut, Y. Sychev, I. De Kouchkovsky, T. Huang, K. Borges, D. Trauner, R. N. Van Gelder, R. H. Kramer, Photochemical Restoration of Visual Responses in Blind Mice. Neuron. 75, 271–282 (2012).

9. L. Laprell, I. Tochitsky, K. Kaur, M. B. Manookin, M. Stein, D. M. Barber, C. Schön, S. Michalakis, M. Biel, R. H. Kramer, M. P. Sumser, D. Trauner, R. N. Van Gelder, Photopharmacological control of bipolar cells restores visual function in blind mice. J. Clin. Invest. 127, 2598–2611 (2017).

10. Kiora Pharmaceuticals, A Phase II Study of Intravitreal KIO-301 in Patients with Late-stage Retinitis Pigmentosa (ABACUS-2) (2024), p. ClinicalTrials.gov ID NCT06628947, (available at https://clinicaltrials.gov/study/NCT06628947?term=Kiora&rank=1#study-overview).

11. X. Yang, D. L. Rode, D. S. Peterka, R. Yuste, S. M. Rothman, Optical control of focal epilepsy in vivo with caged gamma-aminobutyric acid. Ann Neurol. 71, 68–75 (2012).

12. D. Wang, Z. Yu, J. Yan, F. Xue, G. Ren, C. Jiang, W. Wang, Y. Piao, X. Yang, Photolysis of caged-GABA rapidly terminates seizures in vivo: Concentration and light intensity dependence. Front. Neurol. 8, 1–10 (2017).

13. J. Spanoghe, E. Wynendaele, M. Vergaelen, M. De Colvenaer, T. Mariman, K. Vonck, E. Carrette, W. Wadman, E. Craey, L. E. Larsen, M. Sprengers, J. Missinne, S. Van Calenbergh, B. De Spiegeleer, D. De Bundel, I. Smolders, P. Boon, R. Raedt, Photopharmacological activation of adenosine A1 receptor signaling suppresses seizures in a mouse model for temporal lobe epilepsy. J. Control. Release. 381, 113626 (2025).

14. J. M. Sanchez-Sanchez, F. Riefolo, A. Barbero-Castillo, R. Sortino, L. Agnetta, A. Manasanch, C. Matera, M. Bosch, M. Forcella, M. Decker, P. Gorostiza, M. V. Sanchez-Vives, Control of cortical slow oscillations and epileptiform discharges with photoswitchable type 1 muscarinic ligands. PNAS Nexus. 4, 1–12 (2025).

15. E. Beghi, G. Giussani, F. Abd-Allah, J. Abdela, A. Abdelalim, H. N. Abraha, M. G. Adib, S. Agrawal, F. Alahdab, A. Awasthi, Y. Ayele, M. A. Barboza, A. B. Belachew, B. Biadgo, A. Bijani, H. Bitew, F. Carvalho, Y. Chaiah, A. Daryani, H. P. Do, M. Dubey, A. Y. Y. Endries, S. Eskandarieh, A. Faro, F. Farzadfar, S. M. Fereshtehnejad, E. Fernandes, D. O. Fijabi, I. Filip, F. Fischer, A. K. Gebre, A. G. Tsadik, T. G. Gebremichael, K. E. Gezae, M. Ghasemi-Kasman, K. G. Weldegwergs, M. G. Degefa, E. V. Gnedovskaya, T. B. Hagos, A. Haj-Mirzaian, A. Haj-Mirzaian, H. Y. Hassen, S. I. Hay, M. Jakovljevic, A. Kasaeian, T. D. Kassa, Y. S. Khader, I. Khalil, E. A. Khan, J. Khubchandani, A. Kisa, K. J. Krohn, C. Kulkarni, Y. L. Nirayo, M. T. Mackay, M. Majdan, A. Majeed, T. Manhertz, M. M. Mehndiratta, T. Mekonen, H. G. Meles, G. Mengistu, S. Mohammed, M. Naghavi, A. H. Mokdad, G. Mustafa, S. S. N. Irvani, L. H. Nguyen, E. Nichols, M. R. Nixon, F. A. Ogbo, A. T. Olagunju, T. O. Olagunju, M. O. Owolabi, M. R. Phillips, G. D. Pinilla-Monsalve, M. Qorbani, A. Radfar, A. Rafay, V. Rahimi-Movaghar, N. Reinig, P. S. Sachdev, H. Safari, S. Safari, S. Safiri, M. A. Sahraian, A. M. Samy, S. Sarvi, M. Sawhney, M. A. Shaikh, M. Sharif, G. Singh, M. Smith, C. E. I. Szoeke, R. Tabarés-Seisdedos, M. H. Temsah, O. Temsah, M. Tortajada-Girbés, B. X. Tran, A. A. T. Tsegay, I. Ullah, N. Venketasubramanian, R. Westerman, A. S. Winkler, E. M. Yimer, N. Yonemoto, V. L. Feigin, T. Vos, C. J. L. Murray, Global, regional, and national burden of epilepsy, 1990–2016: a systematic analysis for the Global Burden of Disease Study 2016. Lancet Neurol. 18, 357–375 (2019).

16. Y. Du, J. Lin, J. Shen, S. Ding, M. Ye, L. Wang, Y. Wang, X. Wang, N. Xia, R. Zheng, H. Chen, H. Xu, Adverse drug reactions associated with six commonly used antiepileptic drugs in southern China from 2003 to 2015. BMC Pharmacol. Toxicol. 20, 1–8 (2019).

17. E. Craey, F. Hulpia, J. Spanoghe, S. Manzella, L. E. Larsen, M. Sprengers, D. De Bundel, I. Smolders, E. Carrette, J. Delbeke, K. Vonck, P. Boon, S. Van Calenbergh, W. J. Wadman, R. Raedt, Ex Vivo Feedback Control of Neurotransmission Using a Photocaged Adenosine A1 Receptor Agonist. Int. J. Mol. Sci. 23, 1–12 (2022).

18. J. B. Trads, K. Hüll, B. S. Matsuura, L. Laprell, T. Fehrentz, N. Görldt, K. A. Kozek, C. D. Weaver, N. Klöcker, D. M. Barber, D. Trauner, Sign Inversion in Photopharmacology: Incorporation of Cyclic Azobenzenes in Photoswitchable Potassium Channel Blockers and Openers. Angew. Chemie - Int. Ed. 58, 15421–15428 (2019).

19. S. Park, G. Loke, Y. Fink, P. Anikeeva, Flexible fiber-based optoelectronics for neural interfaces. Chem. Soc. Rev. 48, 1826–1852 (2019).

20. M. Beaussier, A. Delbos, A. Maurice-Szamburski, C. Ecoffey, L. Mercadal, Perioperative Use of Intravenous Lidocaine. Drugs. 78, 1229–1246 (2018).

21. Q. Yin, W. Zhang, B. Ke, J. Liu, W. Zhang, Lido-OH, a Hydroxyl Derivative of Lidocaine, Produced a Similar Local Anesthesia Profile as Lidocaine With Reduced Systemic Toxicities. Front. Pharmacol. 12, 1–8 (2021).

22. J. Vuyk, E. Sitsen, M. Reekers, in Miller’s Anesthesia, M. A. Gropper, L. I. Eriksson, L. A. Fleisher, N. H. Cohen, K. Leslie, Eds. (Elsevier, ed. 10th, 2024).

23. B. C. Lang, J. Yang, Y. Wang, Y. Luo, Y. Kang, J. Liu, W. S. Zhang, An improved design of water-soluble propofol prodrugs characterized by rapid onset of action. Anesth. Analg. 118, 745–754 (2014).

24. M. Stein, S. J. Middendorp, V. Carta, E. Pejo, D. E. Raines, S. A. Forman, E. Sigel, D. Trauner, Azo-propofols: Photochromic potentiators of GABAA receptors. Angew. Chemie - Int. Ed. 51, 10500–10504 (2012).

25. I. Tochitsky, Z. Helft, V. Meseguer, R. B. Fletcher, K. A. Vessey, M. Telias, B. Denlinger, J. Malis, E. L. Fletcher, R. H. Kramer, How Azobenzene Photoswitches Restore Visual Responses to the Blind Retina. Neuron. 92, 100–113 (2016).

26. E. Krook-Magnuson, C. Armstrong, M. Oijala, I. Soltesz, On-demand optogenetic control of spontaneous seizures in temporal lobe epilepsy. Nat Commun. 4, 1376 (2013).

27. J. P. Andrews, J. Geng, K. Voitiuk, M. A. T. Elliott, D. Shin, A. Robbins, A. Spaeth, A. Wang, L. Li, D. Solis, M. G. Keefe, J. L. Sevetson, J. A. R. De Jesús, K. C. Donohue, H. H. Larson, D. Ehrlich, K. I. Auguste, S. Salama, V. Sohal, T. Sharf, D. Haussler, C. R. Cadwell, Multimodal evaluation of network activity and optogenetic interventions in human hippocampal slices. Nat. Neurosci. 27, 2487–2499 (2024).

28. R. Chen, F. Gore, Q. A. Nguyen, C. Ramakrishnan, S. Patel, S. H. Kim, M. Raffiee, Y. S. Kim, B. Hsueh, E. Krook-Magnusson, I. Soltesz, K. Deisseroth, Deep brain optogenetics without intracranial surgery. Nat. Biotechnol. 39, 161–164 (2021).

29. A. Pitzschke, B. Lovisa, O. Seydoux, M. Zellweger, M. Pfleiderer, Y. Tardy, G. Wagniéres, Red and NIR light dosimetry in the human deep brain. Phys. Med. Biol. 60, 2921–2937 (2015).

30. T. A. Henderson, L. D. Morries, Near-infrared photonic energy penetration: Can infrared phototherapy effectively reach the human brain? Neuropsychiatr. Dis. Treat. 11, 2191–2208 (2015).

31. S. Wang, S. Szobota, Y. Wang, M. Volgraf, Z. Liu, C. Sun, D. Trauner, E. Y. Isacoff, X. Zhang, All Optical Interface for Parallel, Remote, and Spatiotemporal Control of Neuronal Activity. Nano Lett. 7, 3859–3863 (2007).

32. J. A. Frank, M. Moroni, R. Moshourab, M. Sumser, G. R. Lewin, D. Trauner, Photoswitchable fatty acids enable optical control of TRPV1. Nat. Commun. 6 (2015), doi:10.1038/ncomms8118.

33. X. Rovira, A. Trapero, S. Pittolo, C. Zussy, A. Faucherre, C. Jopling, J. Giraldo, J. P. Pin, P. Gorostiza, C. Goudet, A. Llebaria, OptoGluNAM4.1, a Photoswitchable Allosteric Antagonist for Real-Time Control of mGlu4 Receptor Activity. Cell Chem. Biol. 23, 929–934 (2016).

34. V. A. Gutzeit, A. Acosta-Ruiz, H. Munguba, S. Häfner, A. Landra-Willm, B. Mathes, J. Mony, D. Yarotski, K. Börjesson, C. Liston, G. Sandoz, J. Levitz, J. Broichhagen, A fine-tuned azobenzene for enhanced photopharmacology in vivo. Cell Chem. Biol. 28, 1648–1663.e16 (2021).

35. W. Yang, R. Yuste, In vivo imaging of neural activity. Nat. Methods. 14, 349–359 (2017).

36. V. Venkataramani, D. I. Tanev, C. Strahle, A. Studier-Fischer, L. Fankhauser, T. Kessler, C. Körber, M. Kardorff, M. Ratliff, R. Xie, H. Horstmann, M. Messer, S. P. Paik, J. Knabbe, F. Sahm, F. T. Kurz, A. A. Acikgöz, F. Herrmannsdörfer, A. Agarwal, D. E. Bergles, A. Chalmers, H. Miletic, S. Turcan, C. Mawrin, D. Hänggi, H. K. Liu, W. Wick, F. Winkler, T. Kuner, Glutamatergic synaptic input to glioma cells drives brain tumour progression. Nature. 573, 532–538 (2019).

37. F. Winkler, Neuroscience and oncology: State-of-the-art and new perspectives. Curr. Opin. Neurol. 36, 544–548 (2023).

38. G. K. Bergey, M. J. Morrell, E. M. Mizrahi, A. Goldman, D. King-Stephens, D. Nair, S. Srinivasan, B. Jobst, R. E. Gross, D. C. Shields, G. Barkley, V. Salanova, P. Olejniczak, A. Cole, S. S. Cash, K. Noe, R. Wharen, G. Worrell, A. M. Murro, J. Edwards, M. Duchowny, D. Spencer, M. Smith, E. Geller, R. Gwinn, C. Skidmore, S. Eisenschenk, M. Berg, C. Heck, P. Van Ness, N. Fountain, P. Rutecki, A. Massey, C. O’Donovan, D. Labar, R. B. Duckrow, L. J. Hirsch, T. Courtney, F. T. Sun, C. G. Seale, Long-term treatment with responsive brain stimulation in adults with refractory partial seizures. Neurology. 84, 810–817 (2015).

39. B. N. Lundstrom, G. M. Osman, K. Starnes, N. M. Gregg, H. D. Simpson, Emerging approaches in neurostimulation for epilepsy. Curr. Opin. Neurol. 36, 69–76 (2023).

40. A. D. Mickle, S. M. Won, K. N. Noh, J. Yoon, K. W. Meacham, Y. Xue, L. A. McIlvried, B. A. Copits, V. K. Samineni, K. E. Crawford, D. H. Kim, P. Srivastava, B. H. Kim, S. Min, Y. Shiuan, Y. Yun, M. A. Payne, J. Zhang, H. Jang, Y. Li, H. H. Lai, Y. Huang, S. Il Park, R. W. Gereau, J. A. Rogers, A wireless closed-loop system for optogenetic peripheral neuromodulation. Nature. 565, 361–365 (2019).

41. E. Song, C. H. Chiang, R. Li, X. Jin, J. Zhao, M. Hill, Y. Xia, L. Li, Y. Huang, S. M. Won, K. J. Yu, X. Sheng, H. Fang, M. A. Alam, Y. Huang, J. Viventi, J. K. Chang, J. A. Rogers, Flexible electronic/optoelectronic microsystems with scalable designs for chronic biointegration. Proc. Natl. Acad. Sci. U. S. A. 116, 15398–15406 (2019).

42. S. Il Park, G. Shin, J. G. McCall, R. Al-Hasani, A. Norrisf, L. Xia, D. S. Brenner, K. N. Noh, S. Y. Bang, D. L. Bhatti, K. I. Jang, S. K. Kang, A. D. Mickle, G. Dussor, T. J. Price, R. W. G. Iv, M. R. Bruchas, J. A. Rogers, Stretchable multichannel antennas in soft wireless optoelectronic implants for optogenetics. Proc. Natl. Acad. Sci. U. S. A. 113, E8169–E8177 (2016).

43. A. Canales, X. Jia, U. P. Froriep, R. A. Koppes, C. M. Tringides, J. Selvidge, C. Lu, C. Hou, L. Wei, Y. Fink, P. Anikeeva, Multifunctional fibers for simultaneous optical, electrical and chemical interrogation of neural circuits in vivo. Nat. Biotechnol. 33, 277–284 (2015).

44. H. P. Mattelaer, C. A. Mattelaer, N. Papastavrou, W. Dehaen, P. Herdewijn, Oligonucleotide promoted peptide bond. Chem. Commun. 53, 5013–5016 (2017).

45. V. J. Stella, J. J. Zygmunt, I. G. Georg, M. S. Safadi, Water soluble prodrugs of hindered alcohols or phenols. Patent: WO2000008033, Application No.: WO1999-US17779 (2000)

46. L. P. Smaga, N. W. Pino, G. E. Ibarra, V. Krishnamurthy, J. Chan, A Photoactivatable Formaldehyde Donor with Fluorescence Monitoring Reveals Threshold to Arrest Cell Migration. J. Am. Chem. Soc. 142, 680–684 (2020).

47. G. R. Fulmer, A. J. M. Miller, N. H. Sherden, H. E. Gottlieb, A. Nudelman, B. M. Stoltz, J. E. Bercaw, K. I. Goldberg, NMR Chemical Shifts of Trace Impurities: Common Laboratory Solvents, Organics, and Gases in Deuterated Solvents Relevant to the Organometallic Chemist. Organometallics. 29, 2176–2179 (2010).

48. E. Kandel, J. Schwartz, T. Jessell, S. Siegelbaum, A. J. Hudspeth, Principles of Neural Science, Fifth Edition (McGraw Hill, 2012).

