## Supplemental_Information for "Targeting epilepsy with photopharmacology in human brain tissue"

**Baues et al.**

**<u>Supplementary Figures, Tables, and NMR/MS Spectra</u>**

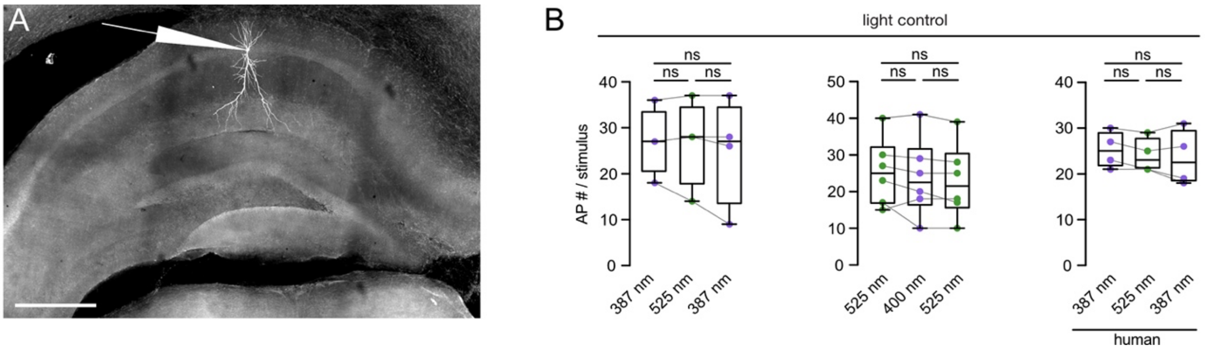


**Suppl. Figure 1: Biocytin staining and QAQ, CQAQ control experiments**
**A)** Biocytin staining of a CA1 neuron in a coronal mouse hippocampal slice co-stained with DAPI. Typical recording location marked by pipette. Scale bar: 500 µm. Blue inset: Magnification of stained neuron. Scale bar: 250 µm. **B)** Quantification of the effect of light-only application (light control) on stimulus-evoked (1 sec, 120 – 600 pA, applied through patch pipette) action potential (AP) firing rate in mouse (left, middle) and human (right) hippocampal neurons. Left (4 cells, 3 mice): 27.00 ± 3.67 (387 nm), 26.75 ± 4.75 (525 nm), 25.00 ± 5.85 ($2^{nd}$ 387 nm). Repeated measures (RM) one-way ANOVA + Tukey's test (F [1.029, 3.086] = 0.826): 387 nm vs. 525 nm p = 0.9783, 525 nm vs. $2^{nd}$ 387 nm p = 0.4130, 387 nm vs. $2^{nd}$ 387 nm p = 0.7079. Center (6 cells, 4 mice): 25.33 ± 3.75 (525 nm), 23.83 ± 4.33 (400 nm), 22.83 ± 4.14 ($2^{nd}$ 525 nm). RM one-way ANOVA + Tukey's test (F (1.192, 5.960) = 2.143): 525 nm vs. 400 nm p = 0.5730, 400 nm vs. $2^{nd}$ 525 nm p = 0.2231, 525 nm vs. $2^{nd}$ 525 nm p = 0.2956. Right (4 cells, 1 patient): 25.25 ± 2.02 (387 nm), 24.00 ± 1.92 (525 nm), 23.50 ± 3.07 ($2^{nd}$ 387 nm). RM one-way ANOVA + Tukey's test (F [1.232, 3.696] = 1.519): 387 nm vs. 525 nm p = 0.1534, 525 nm vs. $2^{nd}$ 387 nm p = 0.9101, 387 nm vs. $2^{nd}$ 387 nm p = 0.4442. For entire fig.: Boxes extend from $25^{th}$ – $75^{th}$ percentiles, inner line represents the median. All ± represent s.e.m.. Depiction of statistical significance: n.s. not significant.

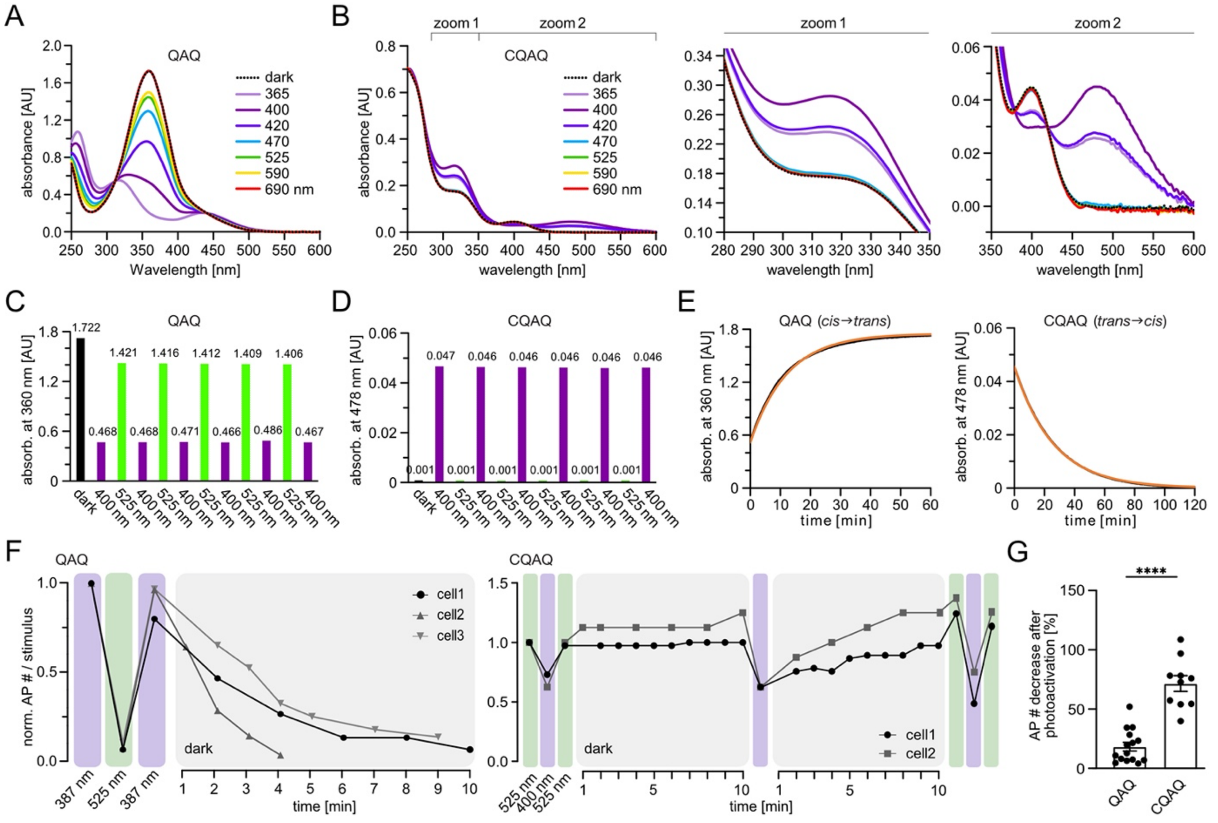


**Suppl. Figure 2: Functional comparison of QAQ and CQAQ**

**A)** Absorption spectra of QAQ (50 µM in PBS buffer, pH 7.4) in the dark-adapted state (thermodynamic equilibrium) and after light irradiation at different wavelengths for 1 min each. Output power was kept equal based on optical power measurements. AU: absorbance unit. **B)** Absorption spectra of CQAQ (50 µM in PBS buffer, pH 7.4) in the dark-adapted state (thermodynamic equilibrium) and after light irradiation at different wavelengths for 1 min each. Output power was kept equal based on optical power measurements. Two zoomed-in spectral sections of the left plot displayed on the right. AU: absorbance unit. **C)** Repeated induced photoswitching of QAQ (50 µM in PBS buffer, pH 7.4) by alternating between λ = 400 nm (purple) and λ = 525 nm (green) irradiation (1 min each cycle) without photodecomposition. Absorbance was measured at the local $\lambda_{max}$ = 360 nm. AU: absorbance unit. **D)** Repeated induced photoswitching of CQAQ (50 µM in PBS buffer, pH 7.4) by alternating between λ = 400 nm (purple) and λ = 525 nm (green) irradiation (1 min each cycle) without photodecomposition. Absorbance was measured at the local $\lambda_{max}$ = 478 nm. AU: absorbance unit. **E)** Thermal relaxation after irradiation (1 min) of QAQ (left) or CQAQ (right; both: 50 µM in PBS buffer, pH 7.4) at 360 nm (QAQ) or 478 nm (CQAQ). Fitted curves are displayed in orange. $t_{1/2}$ (QAQ) = 8.47 min (508 s). $t_{1/2}$ (CQAQ) = 17.9 min (1070 s). AU: absorbance unit. **F)** Representative current clamp recordings in mouse hippocampal neurons (CA1); light-dependent effects on neuronal firing and thermal relaxation dynamics of QAQ ([left] 3 cells, 1 mouse) and CQAQ ([right] 2 cells, 2 mice) in the dark. Note dark-activation of QAQ within minutes, while CQAQ displays higher stability and dark-inactivation. **G)** Quantification of the effect size of QAQ and CQAQ on stimulus-evoked action potential (AP) firing rate in mouse hippocampal neurons (CA1). QAQ (15 cells, 11 mice): 18.57 ± 3.63, CQAQ (10 cells, 5 mice): 71.72 ± 6.60. Unpaired t-test: df = 23; $p < 0.0001$. For entire fig.: All ± represent s.e.m.. Depiction of statistical significance: n.s. not significant, ****$p < 0.0001$.

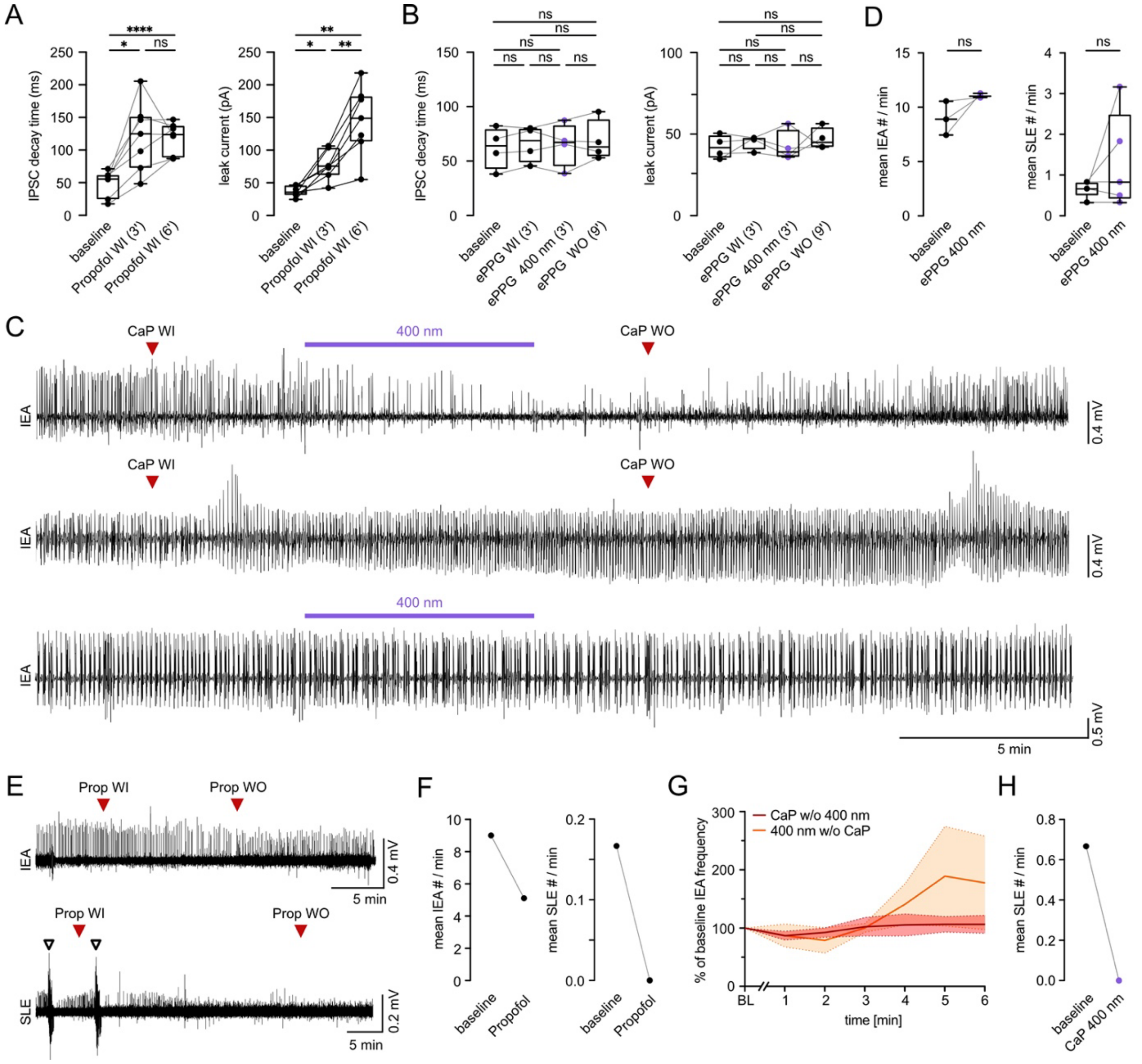


**Suppl. Figure 3: Additional CaP, ePPG and native propofol controls, and analysis of effects of light-activated CaP on SLE in human tissue**

**A)** Quantification of the effect of native propofol application (positive control) on stimulus-evoked (0.2 ms, 100 - 530 μA, applied via bipolar electrode in Schaffer collaterals) inhibitory postsynaptic current (IPSC) decay time (ms) and leak-current (pA) in mouse hippocampal neurons (CA1). Left: IPSC decay time (7 cells, 5 mice), 49.28 ± 7.62 (baseline), 121.96 ± 20.02 (Propofol wash in [WI] 3'), 119.59 ± 8.84 (Propofol WI 6'). Repeated measures (RM) one-way ANOVA + Tukey's test (F [1.040, 6.242] = 20.386): baseline vs. Propofol WI 3' p = 0.0112, Propofol WI 3' vs. Propofol WI 6' p = 0.9997, baseline vs. Propofol WI 6' p < 0.0001. Right: Leak current (7 cells, 5 mice), 36.87 ± 3.06 (baseline), 77.05 ± 8.52 (Propofol WI 3'), 145.58 ± 20.34 (Propofol WI 6'). RM one-way ANOVA + Tukey's test (F [1.176, 7.058] = 28.899): baseline vs. Propofol WI 3' p = 0.0123, Propofol WI 3' vs. Propofol WI 6' p = 0.0059, baseline vs. Propofol WI 6' p = 0.0041. **B)** Quantification of the effect of light and end product photoprotective group (ePPG) application (neg. control) on stimulus-evoked (0.2 ms, 250 - 470 μA, applied via bipolar electrode in Schaffer collaterals) IPSC decay time (ms) and leak-current (pA) in mouse hippocampal neurons (CA1). Left: IPSC decay time (4 cells, 3 mice), 62.00 ± 9.52 (baseline), 65.76 ± 8.28 (ePPG WI), 65.04 ± 10.09 (ePPG 400 nm), 68.55 ± 9.38 (ePPG wash out [WO]). RM one-way ANOVA + Tukey's test (F [1.854, 5.562] = 0.849): baseline vs. ePPG WI p = 0.7503, ePPG WI vs. ePPG 400 nm p > 0.9999, ePPG 400 nm vs. ePPG WO p = 0.9863, baseline vs. ePPG 400 nm p = 0.8498, baseline vs. ePPG WO p = 0.8016, ePPG WI vs. ePPG WO p = 0.9982. Right: Leak current (4 cells, 3 mice), 42.17 ± 3.56 (baseline), 45.14 ± 2.18 (ePPG WI), 42.57 ± 4.75 (ePPG 400 nm), 47.05 ± 3.35 (ePPG WO). RM one-way ANOVA + Tukey's test (F [1.445, 4.335] = 0.473): baseline vs. ePPG WI p = 0.9724, ePPG WI vs. ePPG 400 nm p = 0.9935, ePPG 400 nm vs. ePPG WO p = 0.9949, baseline vs. ePPG 400 nm p > 0.9999, baseline vs. ePPG WO p = 0,9314, ePPG WI vs. ePPG WO p = 0.9990. **C-H)** Interictal epileptiform activity (IEA) and seizure-like episodes (SLE) were evoked by Mg2+-free ACSF solution in both murine (C,D) and human (E-H) cortical slices. **C)** Representative traces displaying extended recording periods in murine slices. Top: Suppression of IEA by CaP light-activation, and recovery of epileptiform activity upon extended wash-out. Middle: control experiment; CaP application without light-activation does not reduce epileptiform activity. Bottom: control experiment; Light-stimulation in the absence of CaP does not reduce epileptiform activity. **D)** Quantification of the effect of light and ePPG application (neg. controls) on IEA # and SLE #. Left: IEA # (3 slices, 2 mice), 8.96 ± 0.90 (baseline), 11.06 ± 0.12 (ePPG 400nm). Paired t-test: df = 2; p = 0.1748. Right: SLE # (5 slices, 2 mice), 0.67 ± 0.09 (baseline), 1.33 ± 0.53 (ePPG 400nm). Paired t-test: df = 4; p = 0.2555. **E)** Representative LFP recording showing the effect of native propofol (positive control) on event number (#) of IEA (top) and SLE (bottom [black arrows]). WI: wash in; WO: wash out; Prop: propofol. **F)** Effect of native propofol (pos. control) on IEA (left) and SLE # (right). Left (1 slice, 1 patient): 9.00 (baseline), 5.11 (propofol). Right (1 slice, 1 patient): 0.17 (baseline), 0.00 (propofol). **G)** Quantification of IEA frequency (normalized to baseline, in %) across 6 min of CaP application without light-activation (4 slices, 3 patients) vs. light-only application (no CaP; 3 slices, 2 patients) negative controls. Multiple unpaired t-tests + Holm-Šídák correction: min1: 86.99 ± 7.63 (CaP w/o 400 nm) vs. 87.17 ± 19.61 (400 nm w/o CaP), p = 0.9932; min2: 92.33 ± 8.04 (CaP w/o 400 nm) vs. 78.92 ± 21.62 (400 nm w/o CaP), p = 0.9031; min3: 102.24 ± 15.87 (CaP w/o 400 nm) vs. 100.30 ± 7.50 (400 nm w/o CaP), p = 0.9932; min4: 105.36 ± 18.88 (CaP w/o 400 nm) vs. 140.66 ± 35.05 (400 nm w/o CaP), p = 0.8920; min5: 106.29 ± 13.38 (CaP w/o 400 nm) vs. 189.04 ± 85.48 (400 nm w/o CaP), p = 0.8920; min6: 106.21 ± 15.37 (CaP w/o 400 nm) vs. 177.53 ± 80.22 (400 nm w/o CaP), p = 0.8920. Dotted lines/shades represent s.e.m. values. BL: baseline. **H)** Effect of light-activated CaP on SLE #. 1 slice, 1 patient: 0.67 (baseline), 0.00 (CaP 400 nm). For entire fig.: Boxes extend from $25^{th}$ – $75^{th}$ percentiles, inner line represents the median. All ± represent s.e.m.. Depiction of statistical significance: n.s. not significant, *p < 0.05, **p < 0.01, ***p < 0.001, ****p < 0.0001.

#### Supplementary Tables

| Age (yrs), Sex | Pathology | Type of epilepsy | Type of surgery | Drug | Type of exp. |
|---|---|---|---|---|---|
| 26, M | MRI-negative | TLE | 2/3 temporal lobectomy | QAQ | light activation |
| 41, M | Low grade glioma | TLE | Anterior temporal lobectomy + AHE | QAQ, and no drug | light activation, and light only (neg. control) |
| 63, M | Pituitary macroadenoma | - | Resection | CQAQ | light activation |
| 48, F | Hippocampal sclerosis | TLE | sAHE | QAQ | light activation |
| 54, M | Hippocampal sclerosis | TLE | 2/3 temporal lobectomy | QAQ | light activation |
| 60, F | Hippocampal sclerosis | TLE | sAHE | QAQ | light activation |
| 29, M | Hippocampal sclerosis | TLE | sAHE | CQAQ | light activation |

**Suppl. Table 1: Clinical data for QAQ, CQAQ experiments shown in Fig. 2 D and Suppl. Fig. 1 B**
MRI = magnetic resonance imaging, TLE = temporal lobe epilepsy,
AHE = amygdalahippocampectomy, sAHE = selective amygdalahippocampectomy

| Thermodynamically stable form | $\lambda$ = 400 nm [a] | $\lambda$ = 525 nm [b] |
|---|---|---|
| *trans*-QAQ | 62 % (*cis*) | 96 % (*trans*) |
| *cis*-CQAQ | 48 % (*trans*) | 98 % (*cis*) |

**Suppl. Table 2: Switching properties of QAQ / CQAQ at 1 mM concentration, room temperature**
[a] % switched after irradiation at $\lambda$ = 400 nm for 10 min
[b] % thermodynamically stable form after irradiation of UV-switched samples at $\lambda$ = 525 nm for 10 min

| Age (yrs), Sex | Pathology | Type of epilepsy | Type of surgery | Drug | Type of exp. |
|---|---|---|---|---|---|
| 43, M | Low grade ganglioglioma | TLE | Temporolateral lesionectomy | Native propofol | no light (pos. control) |
| 19, M | Hippocampal sclerosis | TLE | sAHE | No drug | light only (neg. control) |
| 28, M | MRI-negative | TLE | Temporobasal resection | No drug | light only (neg. control) |
| 57, M | Glioblastoma | - | Anterior temporal lobectomy + AHE | CaP | light activation |
| 37, F | Meningoencephalocele | TLE | Temporal pole resection sparing AMD/HPC | CaP | light activation |
| 53, M | Glioblastoma | TLE | Anterior temporal lobectomy + AHE | CaP | light activation |
| 77, M | Glioblastoma | TLE | Anterior temporal lobectomy | CaP | no light activation (neg. control) |
| 25, M | Suspected FCD | Focal E. | Extended lesionectomy | CaP | light activation |
| 38, M | Cavernoma | TLE | Temporal pole resection sparing AMD/HPC | CaP | no light activation (neg. control) |
| 30, M | Oligodendroglioma | Focal E. | Temporoparietal lesionectomy | CaP | no light activation (neg. control) |

**Suppl. Table 3: Clinical data for CaP experiments shown in Fig. 3 E-J and Suppl. Fig. 3 E-H.**
TLE = temporal lobe epilepsy, sAHE = selective amygdalahippocampectomy, MRI = magnetic resonance imaging, AHE = amygdalahippocampectomy, AMD = amygdala, HPC = hippocampus, FCD = focal cortical dysplasia.

### $^{1}$H-NMR, $^{13}$C-NMR, HRMS and LCMS Spectra

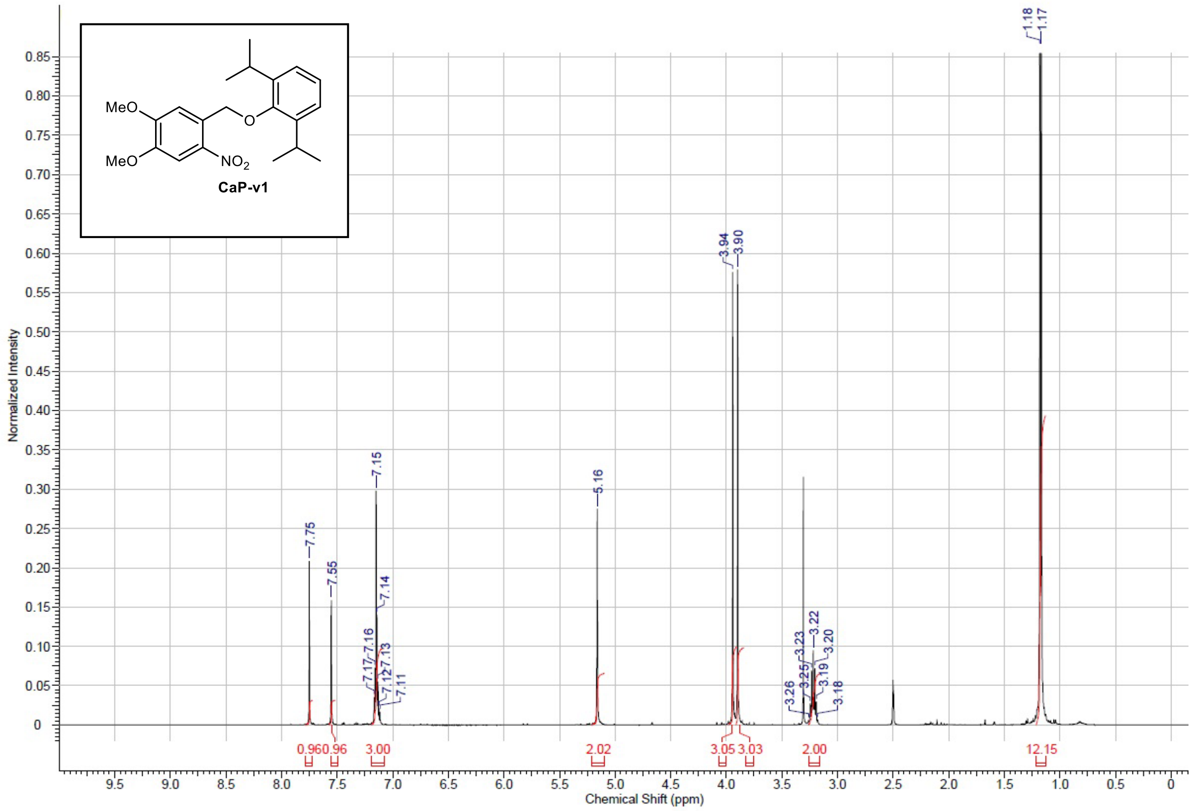


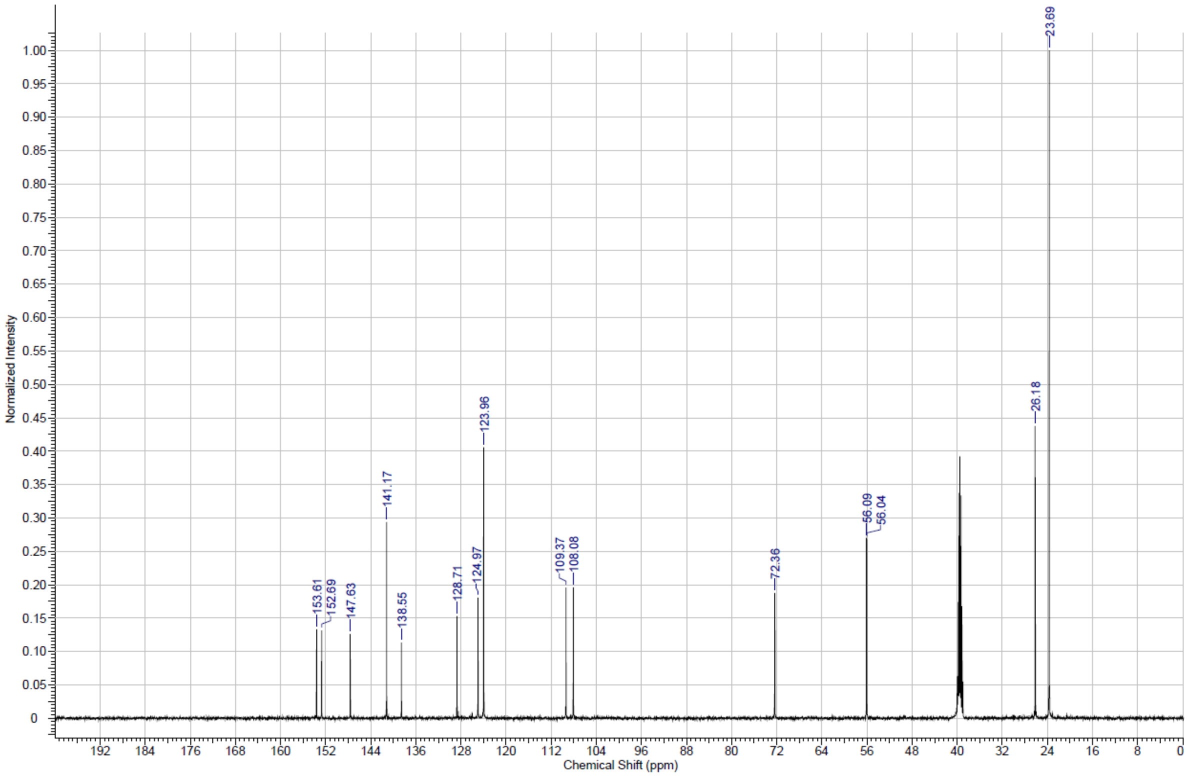

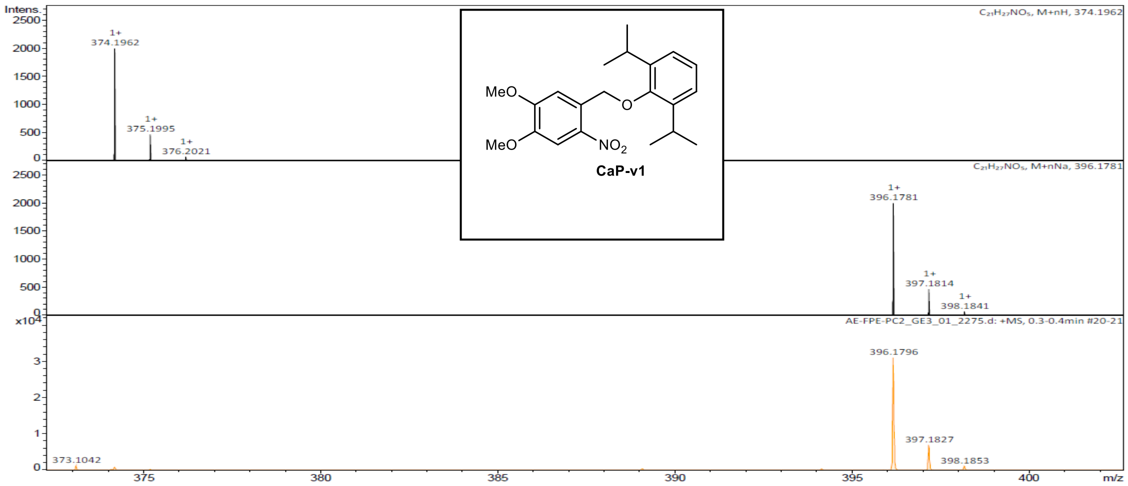


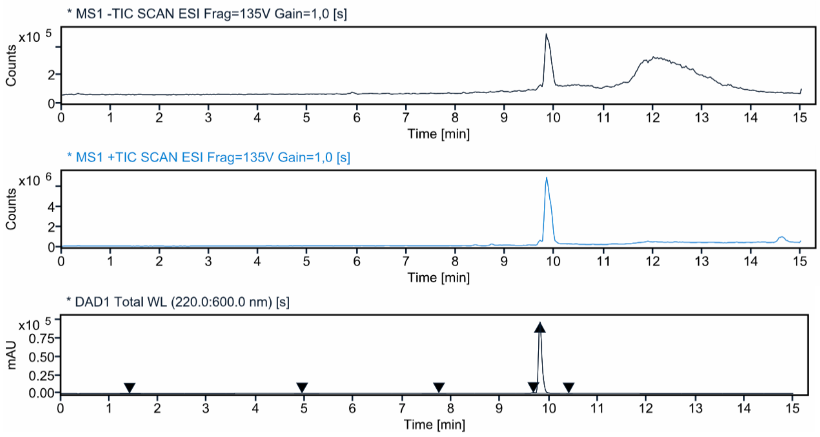


**Signal:** * DAD1 Total WL (220.0:600.0 nm) [s]

| RT [min] | Peak MS Base Peak m/z | Area | Area% | Max Peak% | Height |
|---|---|---|---|---|---|
| 1.414 | | 772.1036 | 0.1586 | 0.160 | 118.215 |
| 4.944 | | 232.2845 | 0.0477 | 0.048 | 107.428 |
| 7.744 | | 407.2176 | 0.0836 | 0.084 | 62.720 |
| 9.679 | | 3083.5172 | 0.6333 | 0.639 | 717.359 |
| 9.812 | | 482255.4483 | 99.0393 | 100.000 | 97562.485 |
| 10.401 | | 183.0686 | 0.0376 | 0.038 | 48.987 |
| | Sum | 486933.6398 | | | |

**Signal:** * MS1 +TIC SCAN ESI Frag=135V Gain=1,0 [s]

| RT [min] | Peak MS Base Peak m/z | Area | Area% | Max Peak% | Height |
|---|---|---|---|---|---|
| 9.845 | 391.300 | 50745417.6349 | 100.0000 | 100.000 | 6373852.558 |
| | Sum | 50745417.6349 | | | |

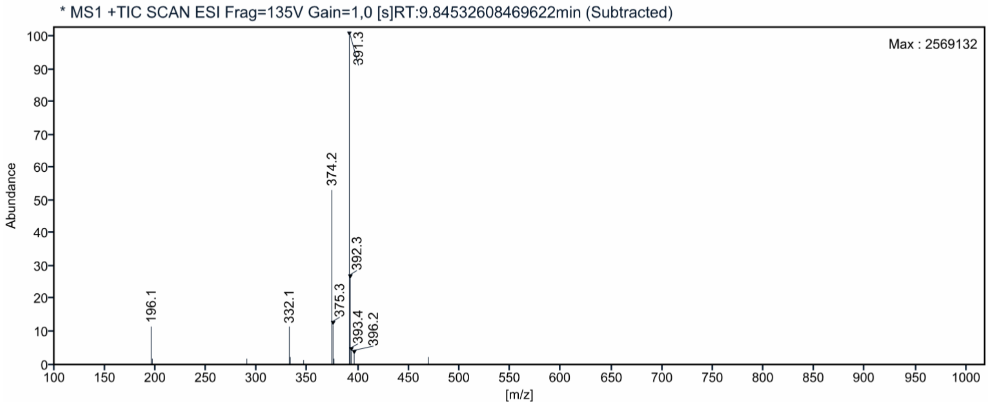

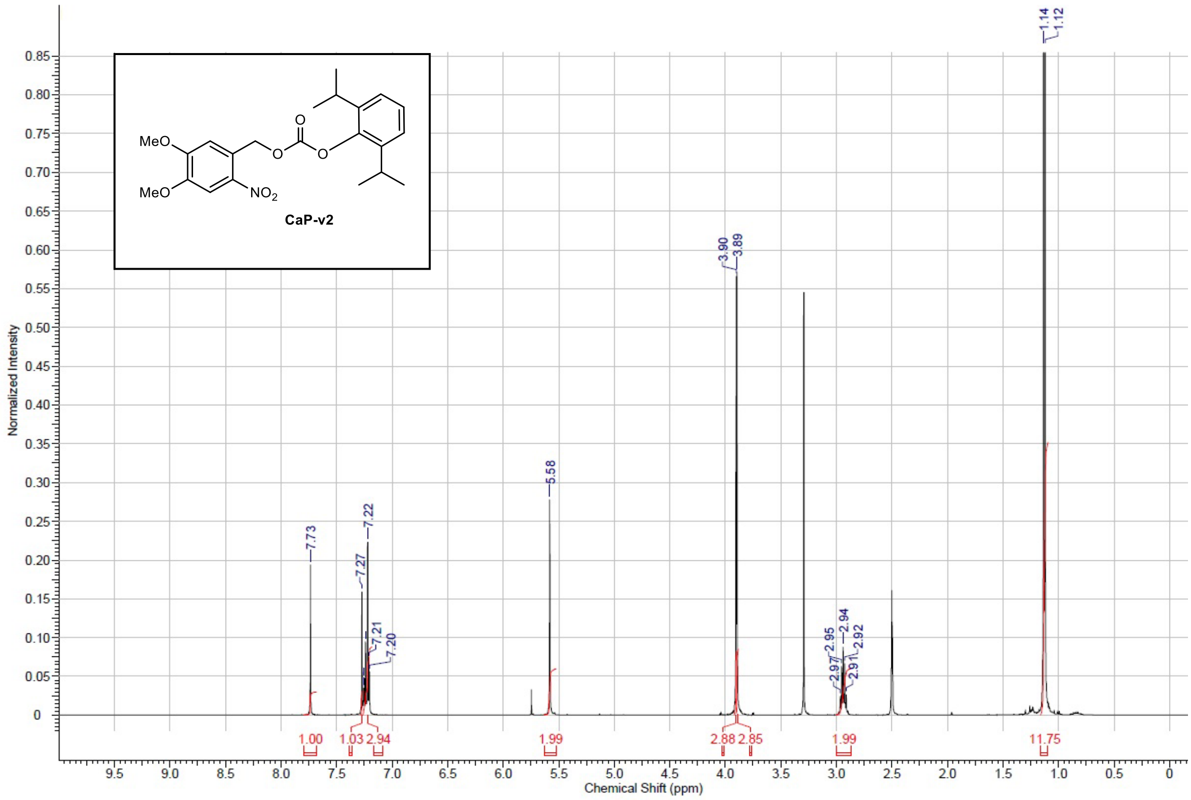
MeO
MeO
O
O
O
NO2
CaP-v2
1.14
1.12
3.90
3.89
5.58
7.73
7.27
7.22
7.21
7.20
2.97
2.95
2.94
2.92
2.91
1.00
1.03
2.94
1.99
2.88
2.85
1.99
11.75
Normalized Intensity
Chemical Shift (ppm)

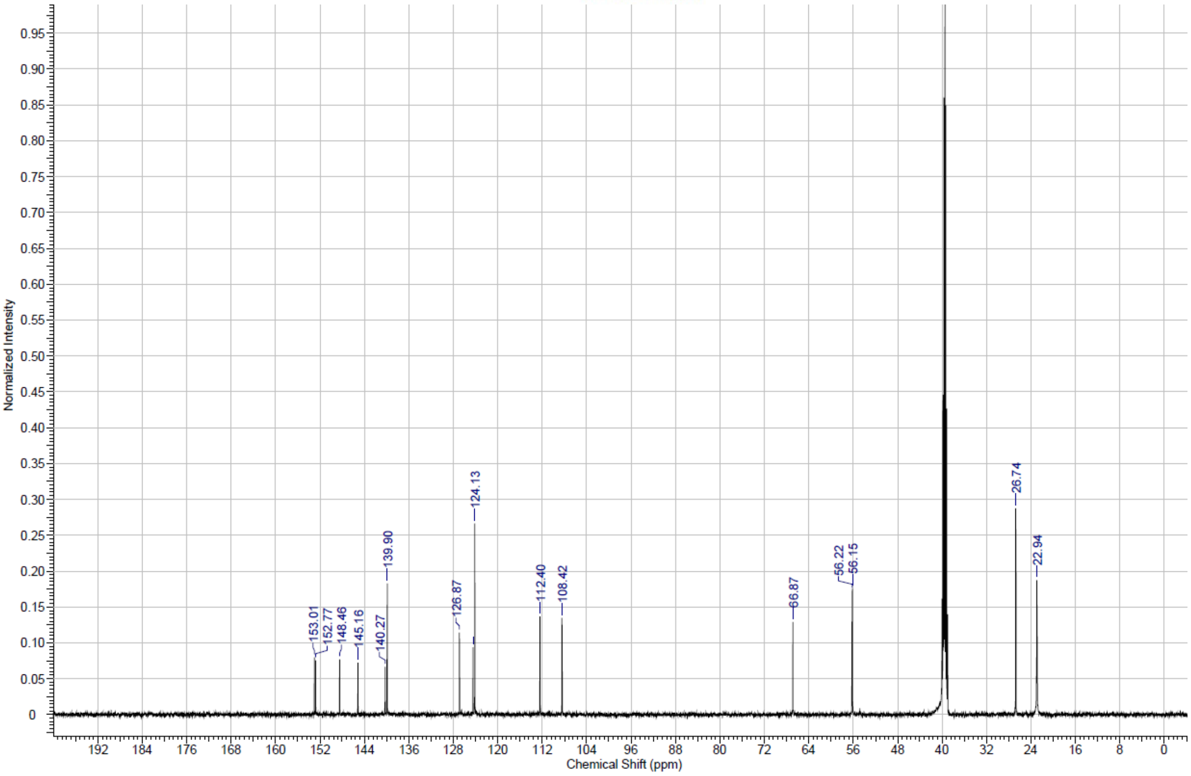
153.01
152.77
148.46
145.16
140.27
139.90
126.87
124.13
112.40
108.42
66.87
56.22
56.15
26.74
22.94
Normalized Intensity
Chemical Shift (ppm)

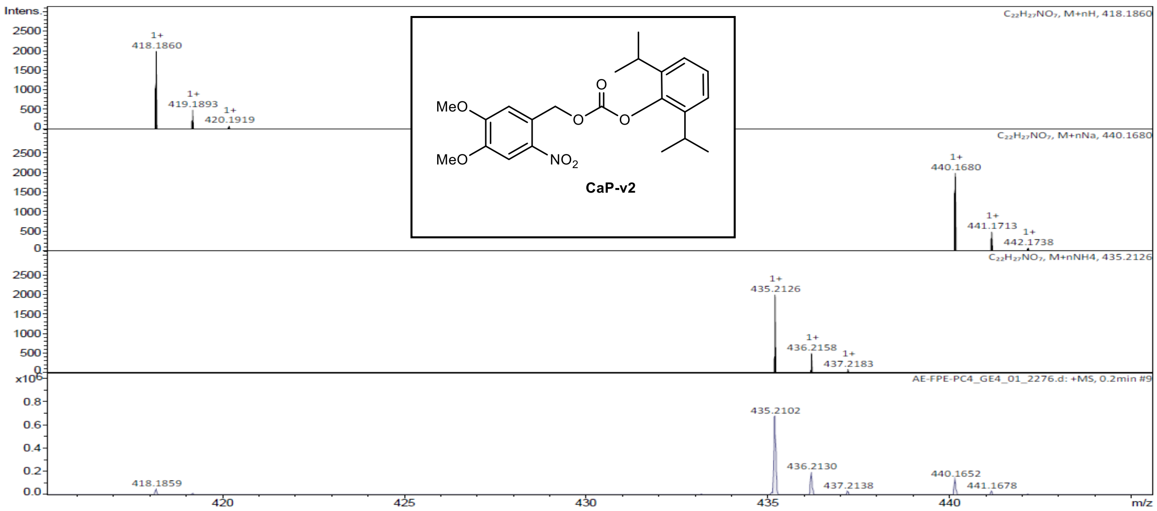


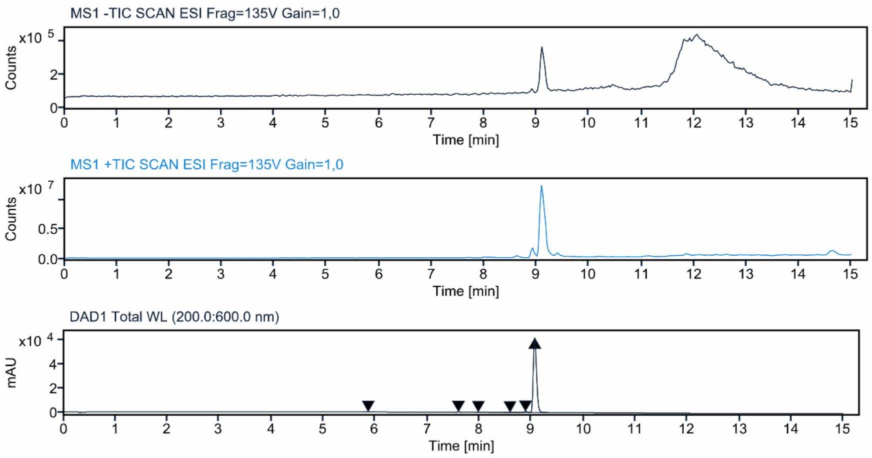


**Signal:** DAD1 Total WL (200.0:600.0 nm)

| RT [min] | Peak MS Base Peak m/z | Area | Area% | Max Peak% | Height |
|---|---|---|---|---|---|
| 5.862 | | 203.9312 | 0.0760 | 0.078 | 54.454 |
| 7.597 | | 805.6600 | 0.3004 | 0.306 | 219.739 |
| 7.980 | | 399.8873 | 0.1491 | 0.152 | 95.141 |
| 8.589 | | 341.4848 | 0.1273 | 0.130 | 58.718 |
| 8.886 | | 3352.8319 | 1.2500 | 1.274 | 818.781 |
| 9.065 | | 263125.4789 | 98.0972 | 100.000 | 62480.827 |
| | Sum | 268229.2740 | | | |

**Signal:** MS1 +TIC SCAN ESI Frag=135V Gain=1,0

| RT [min] | Peak MS Base Peak m/z | Area | Area% | Max Peak% | Height |
|---|---|---|---|---|---|
| 9.098 | 435.400 | 85806622.0303 | 100.0000 | 100.000 | 12410757.067 |
| | Sum | 85806622.0303 | | | |

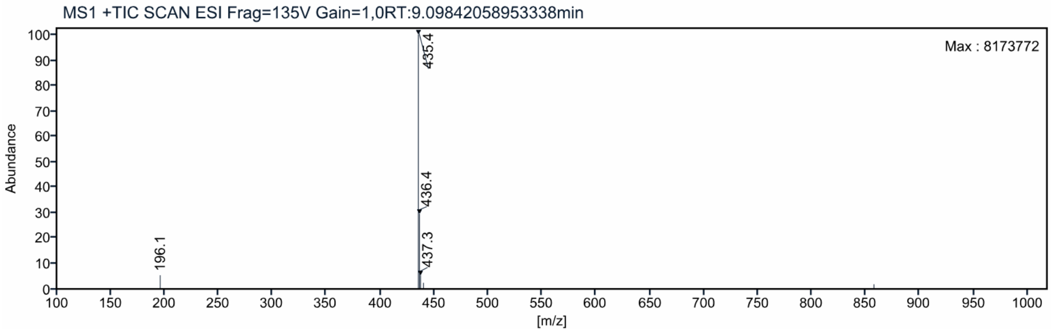

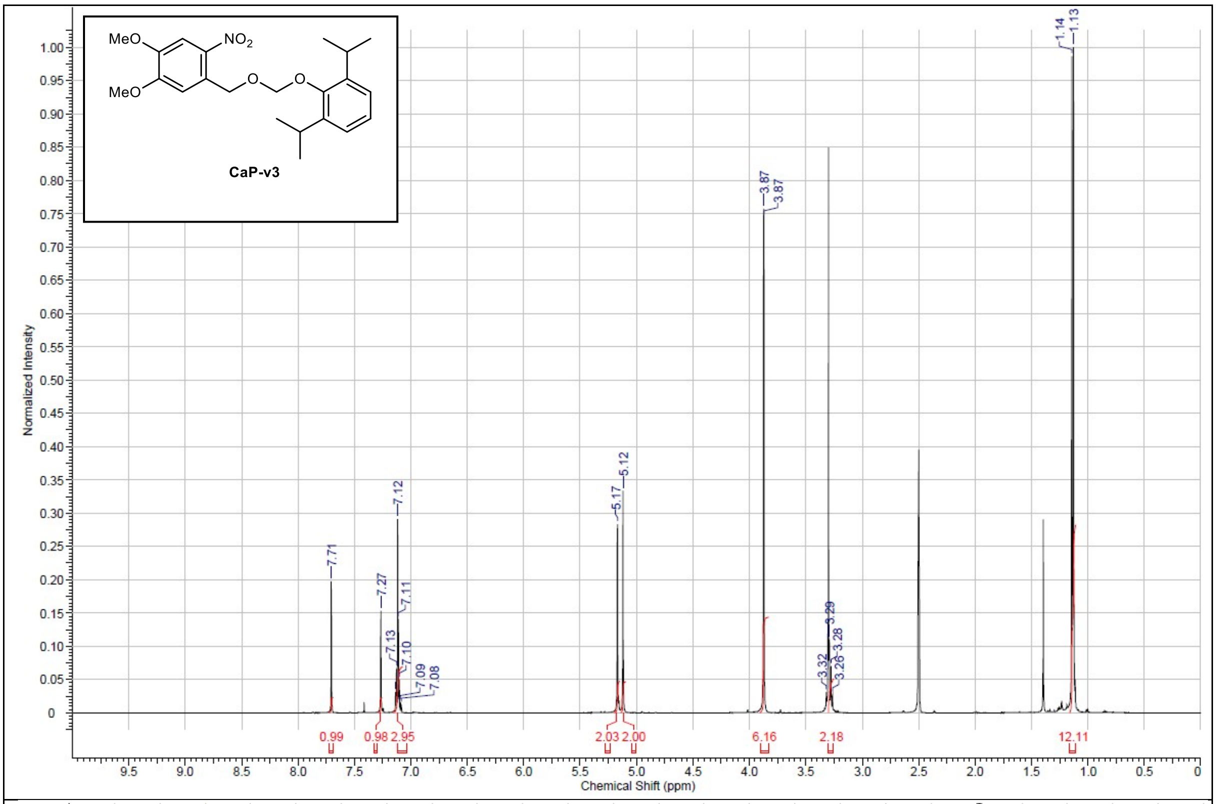
MeO
NO2
MeO
O
O
CaP-v3
Normalized Intensity
Chemical Shift (ppm)
7.71
7.27
7.13
7.12
7.11
7.10
7.09
7.08
5.17
5.12
3.87
3.87
3.32
3.29
3.28
3.26
1.14
1.13
0.99
0.98
2.95
2.03
2.00
6.16
2.18
12.11

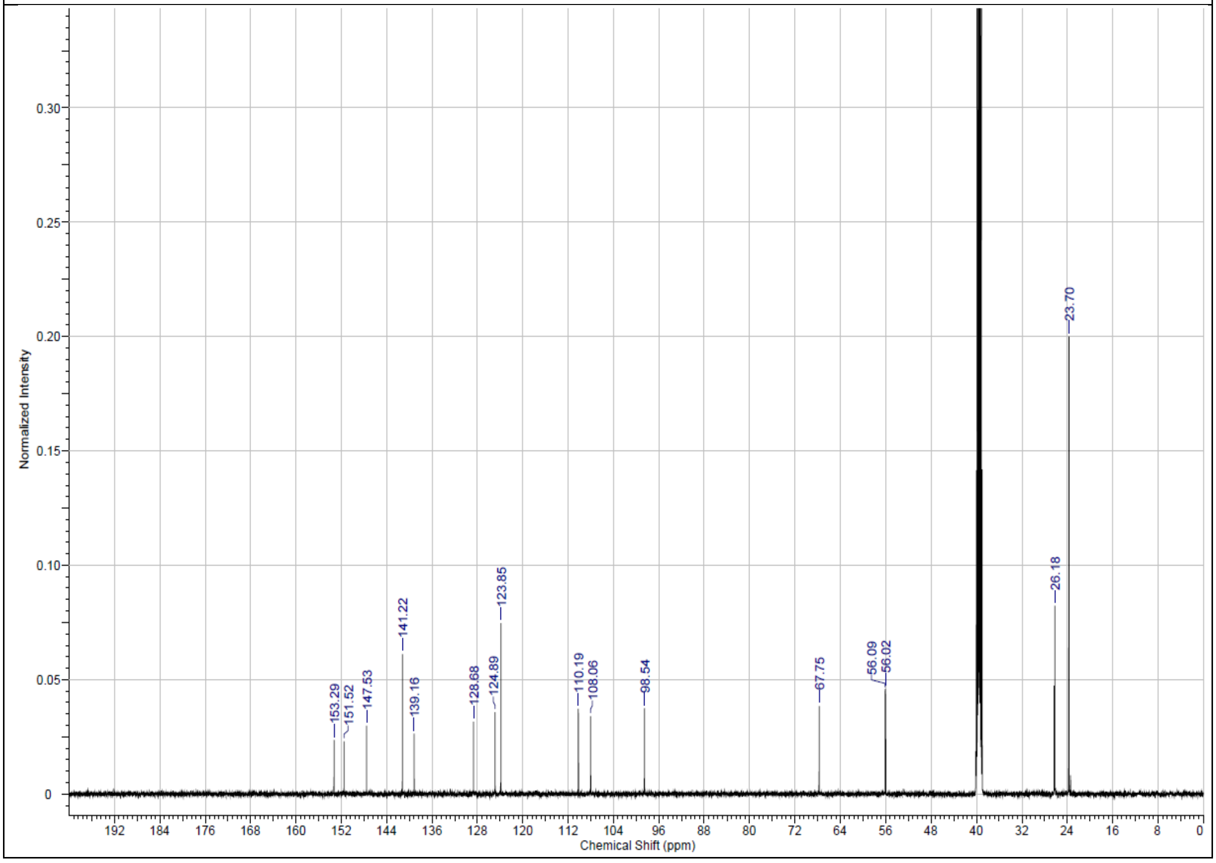
Normalized Intensity
Chemical Shift (ppm)
153.29
151.52
147.53
141.22
139.16
128.68
124.89
123.85
110.19
108.06
98.54
67.75
56.09
56.02
26.18
23.70

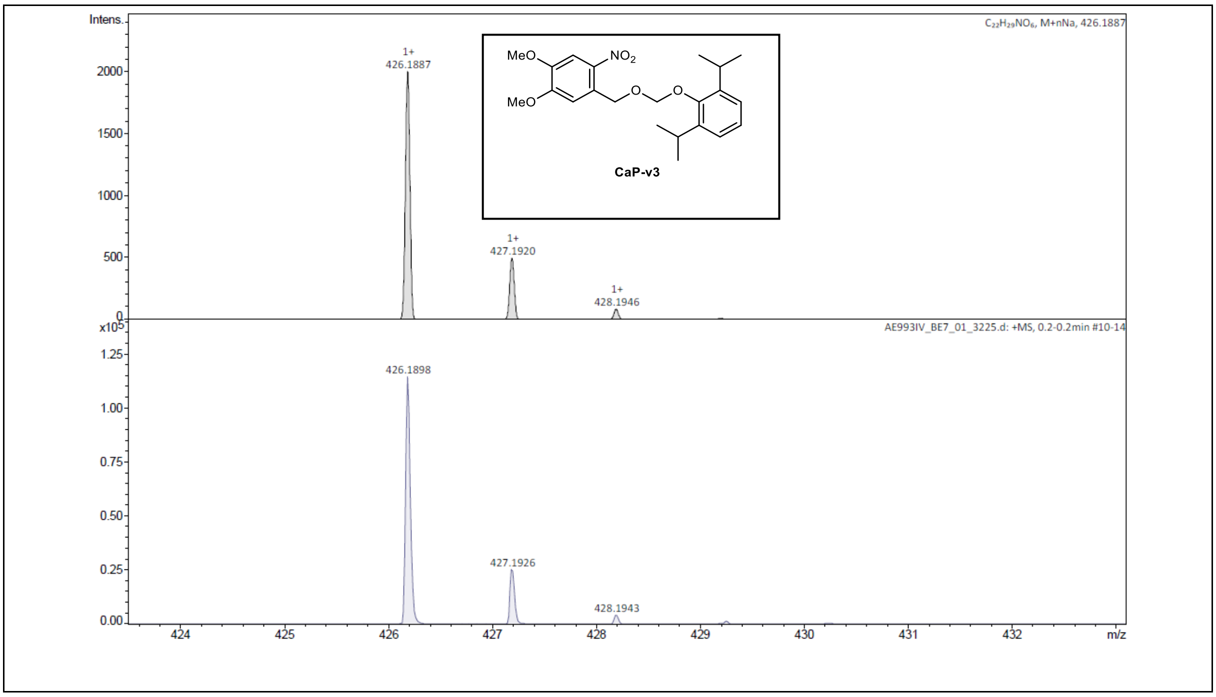


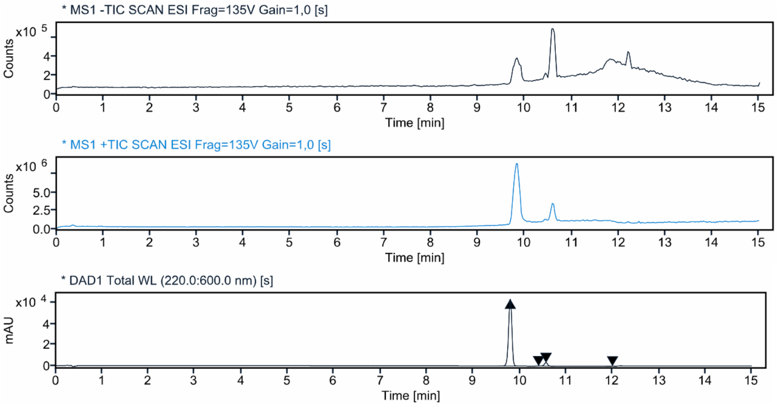


**Signal:** * DAD1 Total WL (220.0:600.0 nm) [s]

| RT [min] | Peak MS Base Peak m/z | Area | Area% | Max Peak% | Height |
|---|---|---|---|---|---|
| 9.797 | 372.500 | 317295.9399 | 95.3675 | 100.000 | 64473.499 |
| 10.406 | | 332.9699 | 0.1001 | 0.105 | 94.586 |
| 10.566 | | 14980.1532 | 4.5025 | 4.721 | 3278.325 |
| 12.000 | | 99.4848 | 0.0299 | 0.031 | 26.255 |
| | Sum | 332708.5477 | | | |

**Signal:** * MS1 +TIC SCAN ESI Frag=135V Gain=1,0 [s]

| RT [min] | Peak MS Base Peak m/z | Area | Area% | Max Peak% | Height |
|---|---|---|---|---|---|
| 9.832 | 372.500 | 75477603.1875 | 100.0000 | 100.000 | 8416232.313 |
| | Sum | 75477603.1875 | | | |

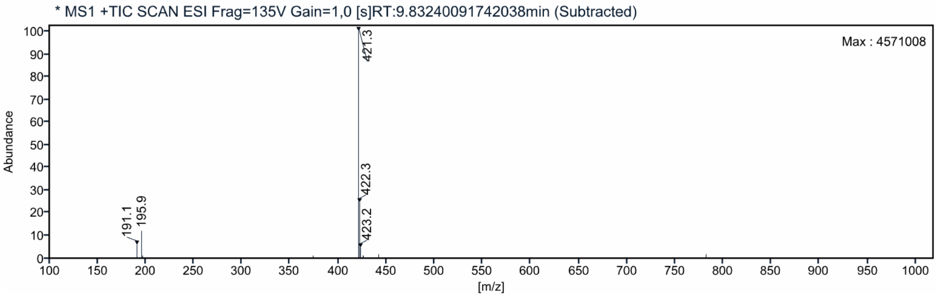

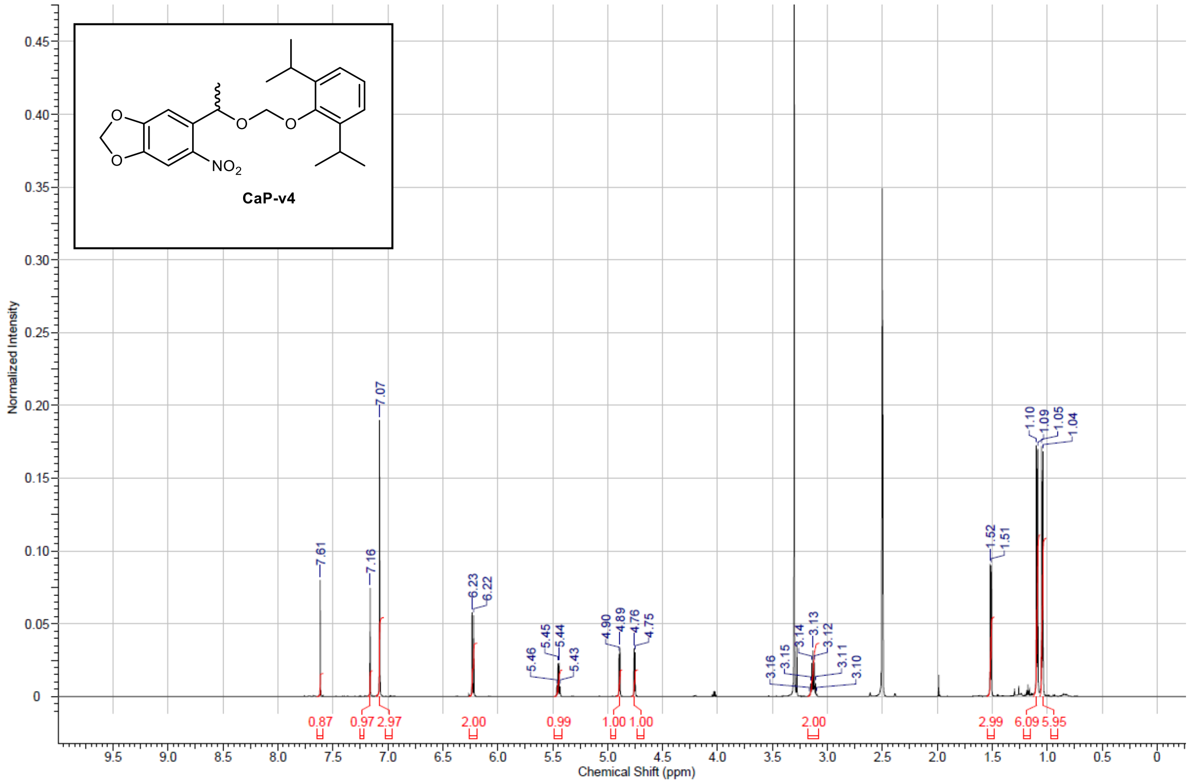

CaP-v4
NO2
O
Normalized Intensity
Chemical Shift (ppm)
7.61
7.16
7.07
6.23
6.22
5.46
5.45
5.44
5.43
4.90
4.89
4.76
4.75
3.16
3.15
3.14
3.13
3.12
3.11
3.10
1.52
1.51
1.10
1.09
1.05
1.04
0.87
0.97
2.97
2.00
0.99
1.00
1.00
2.00
2.99
6.09
5.95


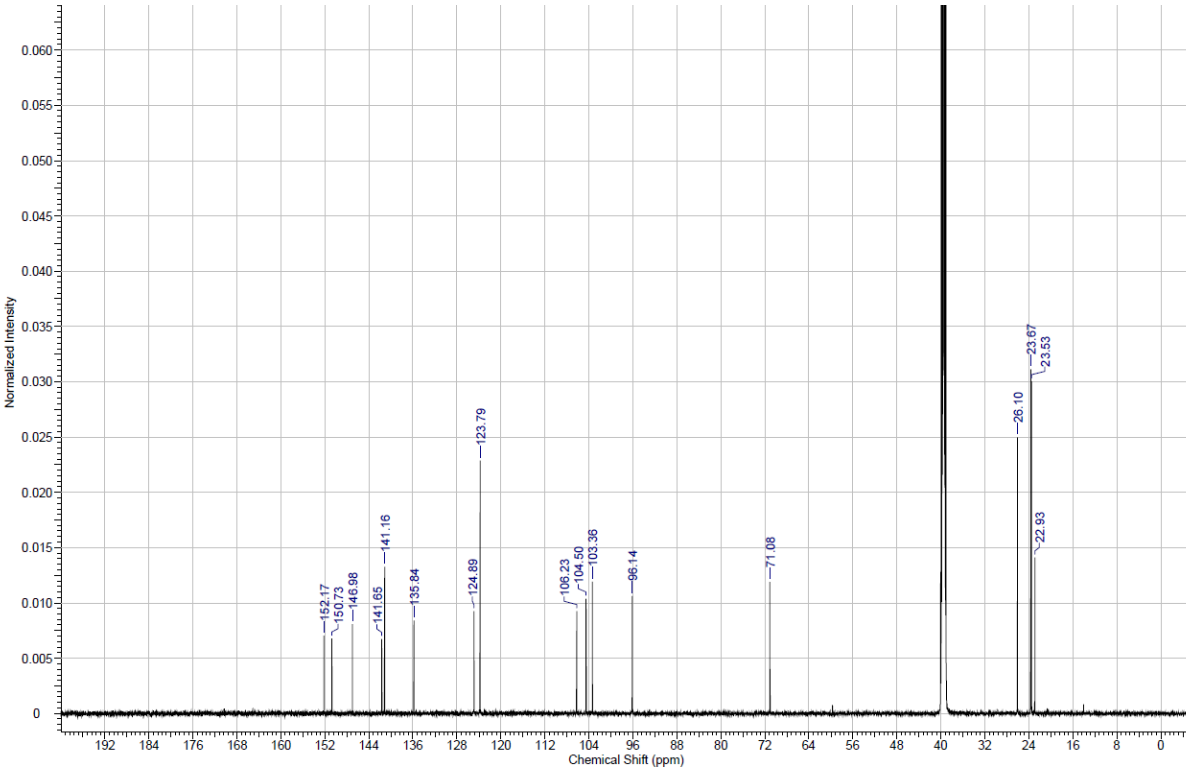

Normalized Intensity
Chemical Shift (ppm)
152.17
150.73
146.98
141.65
141.16
135.84
124.89
123.79
106.23
104.50
103.36
96.14
71.08
26.10
23.67
23.53
22.93

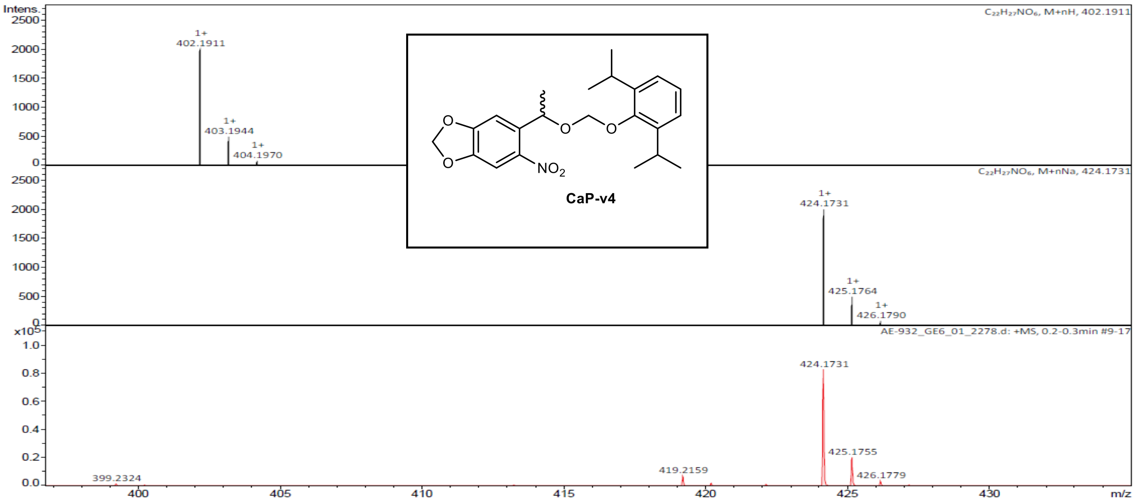


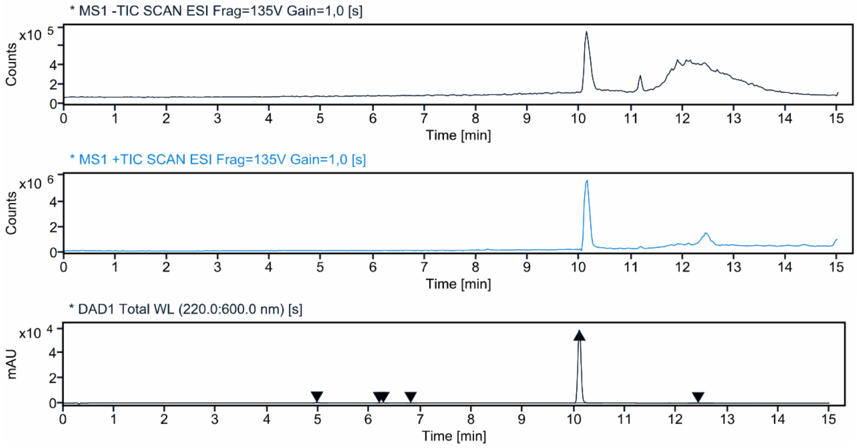


**Signal:** * DAD1 Total WL (220.0:600.0 nm) [s]

| RT [min] | Peak MS Base Peak m/z | Area | Area% | Max Peak% | Height |
|---|---|---|---|---|---|
| 4.971 | | 1896.1097 | 0.7271 | 0.738 | 403.294 |
| 6.189 | | 71.5342 | 0.0274 | 0.028 | 26.439 |
| 6.270 | | 689.2451 | 0.2643 | 0.268 | 155.658 |
| 6.805 | | 178.7333 | 0.0685 | 0.070 | 40.034 |
| 10.106 | | 256968.2820 | 98.5401 | 100.000 | 59148.240 |
| 12.430 | | 971.5250 | 0.3726 | 0.378 | 86.491 |
| | Sum | 260775.4294 | | | |

**Signal:** * MS1 +TIC SCAN ESI Frag=135V Gain=1,0 [s]

| RT [min] | Peak MS Base Peak m/z | Area | Area% | Max Peak% | Height |
|---|---|---|---|---|---|
| 10.140 | 391.300 | 45587115.0655 | 100.0000 | 100.000 | 5596398.607 |
| | Sum | 45587115.0655 | | | |

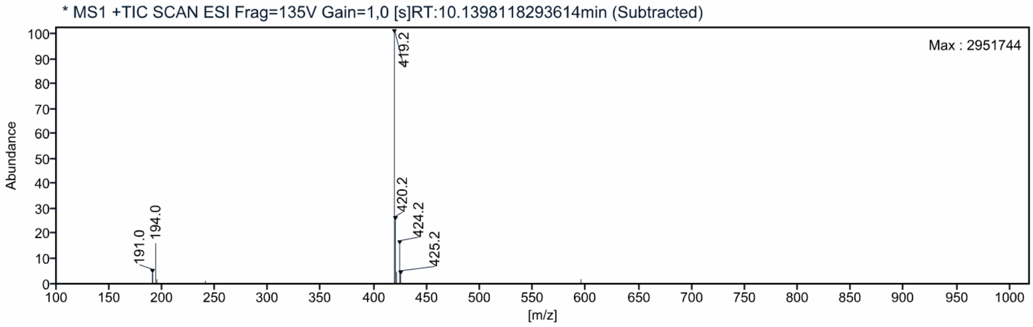

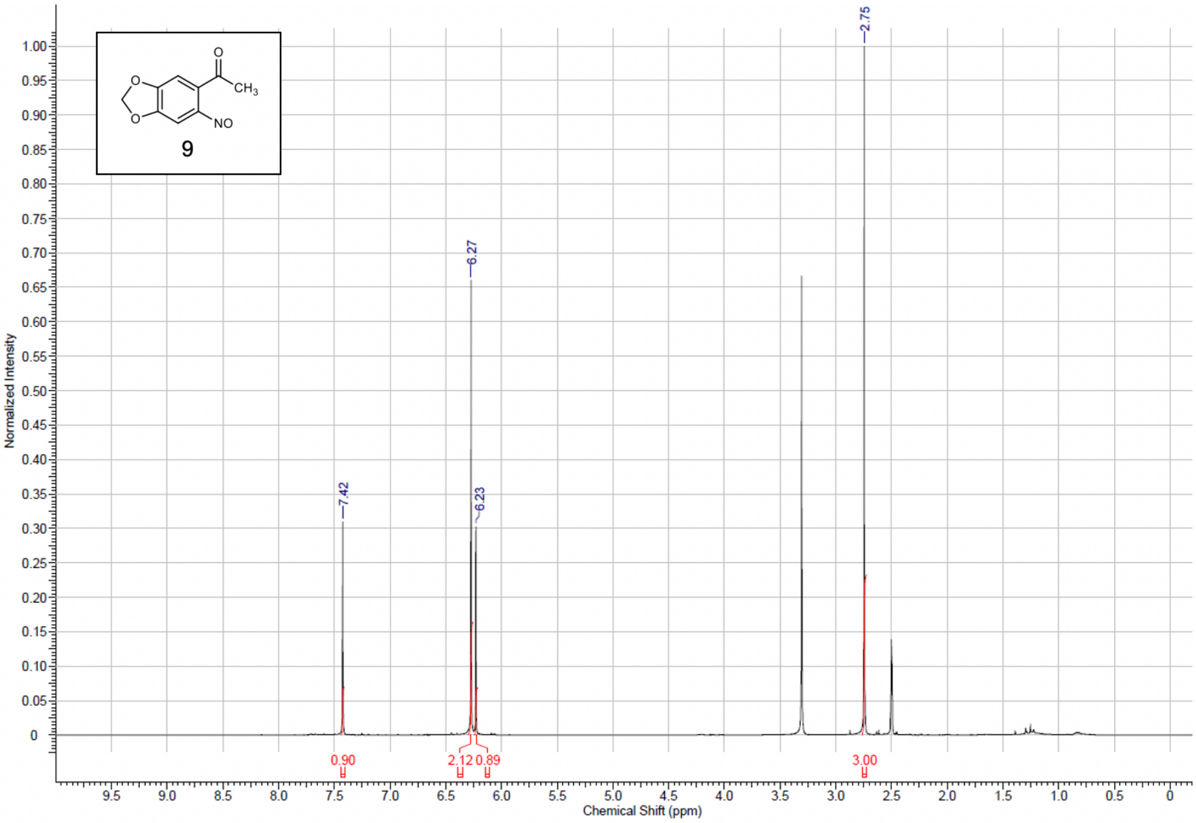
CH3
NO
9
7.42
6.27
6.23
2.75
0.90
2.12
0.89
3.00
Normalized Intensity
Chemical Shift (ppm)

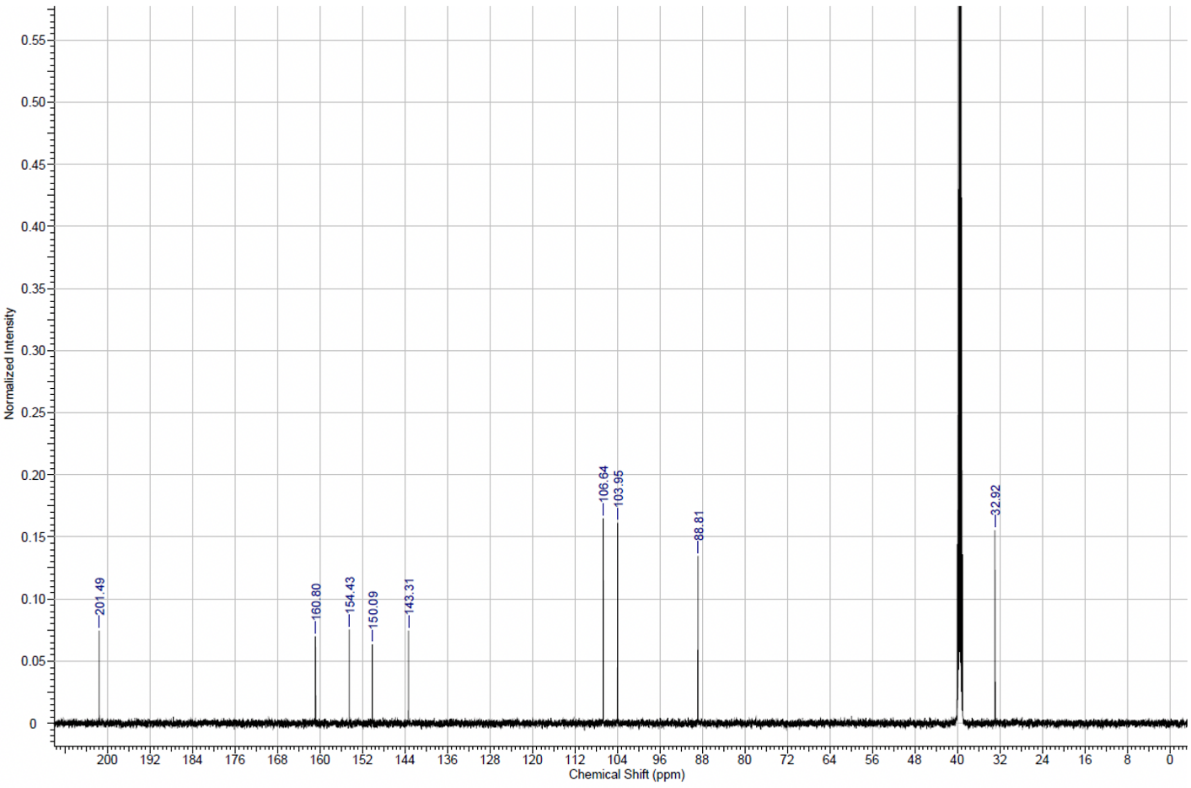
201.49
160.80
154.43
150.09
143.31
106.64
103.95
88.81
32.92
Normalized Intensity
Chemical Shift (ppm)

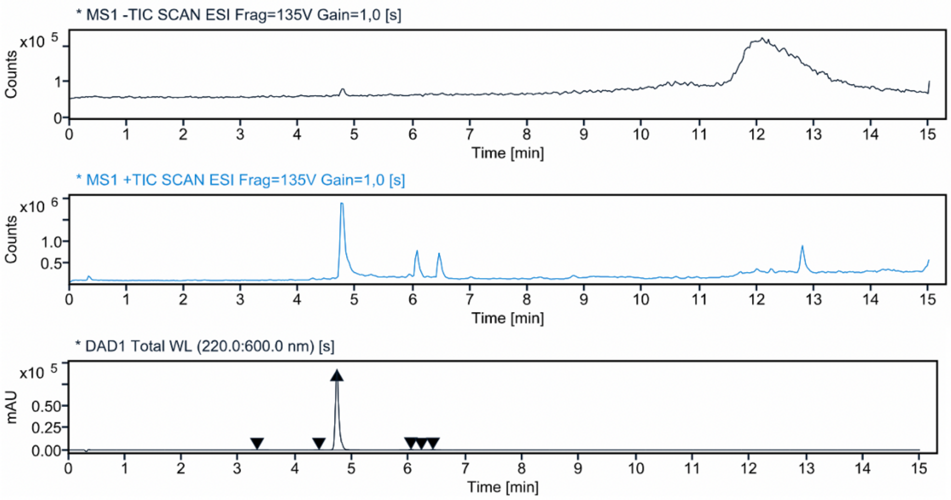


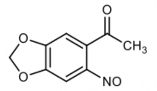


9

**Signal:** * DAD1 Total WL (220.0:600.0 nm) [s]

| RT [min] | Peak MS Base Peak m/z | Area | Area% | Max Peak% | Height |
|---|---|---|---|---|---|
| 3.324 | | 612.0806 | 0.1453 | 0.147 | 119.054 |
| 4.415 | | 353.5748 | 0.0839 | 0.085 | 101.013 |
| 4.729 | | 415753.8275 | 98.6638 | 100.000 | 92854.564 |
| 6.033 | | 2580.5020 | 0.6124 | 0.621 | 604.299 |
| 6.218 | | 218.5854 | 0.0519 | 0.053 | 76.872 |
| 6.422 | | 1865.8006 | 0.4428 | 0.449 | 478.408 |
| | Sum | 421384.3709 | | | |

**Signal:** * MS1 +TIC SCAN ESI Frag=135V Gain=1,0 [s]

| RT [min] | Peak MS Base Peak m/z | Area | Area% | Max Peak% | Height |
|---|---|---|---|---|---|
| 4.760 | 194.000 | 11127450.6602 | 100.0000 | 100.000 | 1911372.696 |
| | Sum | 11127450.6602 | | | |

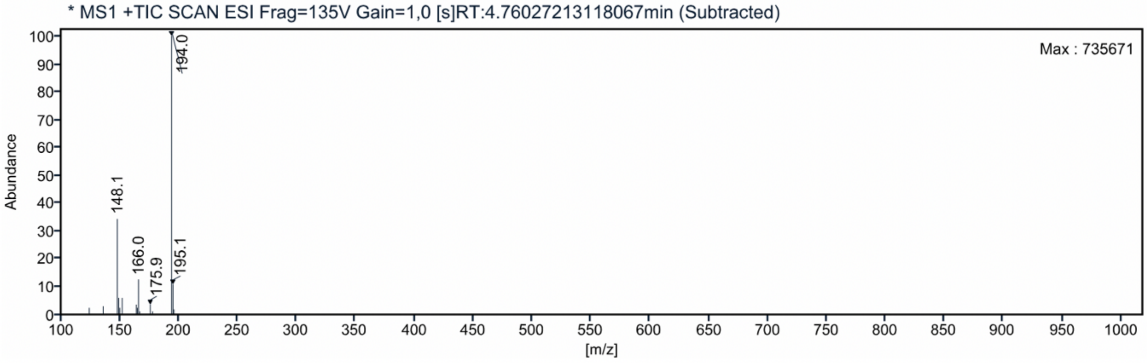
